# Disruption of the PGE_2_ synthesis / response pathway restrains atherogenesis in programmed cell death-1 (Pd-1) deficient hyperlipidemic mice

**DOI:** 10.1101/2024.07.02.601762

**Authors:** Emanuela Ricciotti, Soon Yew Tang, Antonijo Mrčela, Ujjalkumar S. Das, Ronan Lordan, Robin Joshi, Soumita Ghosh, Justin Aoyama, Ryan McConnell, Jianing Yang, Gregory R. Grant, Garret A. FitzGerald

## Abstract

Immune checkpoint inhibitors (ICIs) that target programmed cell death 1 (PD-1) have revolutionized cancer treatment by enabling the restoration of suppressed T-cell cytotoxic responses. However, resistance to single-agent ICIs limits their clinical utility. Combinatorial strategies enhance their antitumor effects, but may also enhance the risk of immune related adverse effects of ICIs.

Prostaglandin (PG) E**_2_**, formed by the sequential action of the cyclooxygenase (COX) and microsomal PGE synthase (mPGES-1) enzymes, acting via its E prostanoid (EP) receptors, EPr2 and EPr4, promotes lymphocyte exhaustion, revealing an additional target for ICIs. Thus, COX inhibitors and EPr4 antagonists are currently being combined with ICIs potentially to enhance antitumor efficacy in clinical trials. However, given the cardiovascular (CV) toxicity of COX inhibitors, such combinations may increase the risk particularly of CV AEs. Here, we compared the impact of distinct approaches to disruption of the PGE**_2_** synthesis /response pathway – global or myeloid cell specific depletion of mPges-1 or global depletion of Epr4 – on the accelerated atherogenesis in Pd-1 deficient hyperlipidemic (Ldlr^-/-^) mice.

All strategies restrained the atherogenesis. While depletion of mPGES-1 suppresses PGE**_2_** biosynthesis, reflected by its major urinary metabolite, PGE**_2_** biosynthesis was increased in mice lacking EPr4, consistent with enhanced expression of aortic Cox-1 and mPges-1. Deletions of mPges-1 and Epr4 differed in their effects on immune cell populations in atherosclerotic plaques; the former reduced neutrophil infiltration, while the latter restrained macrophages and increased the infiltration of T-cells. Consistent with these findings, chemotaxis by bone-marrow derived macrophages from Epr4^-/-^ mice was impaired. Epr4 depletion also resulted in extramedullary lymphoid hematopoiesis and inhibition of lipoprotein lipase activity (LPL) with coincident spelenomegaly, leukocytosis and dyslipidemia. Targeting either mPGES-1 or EPr4 may restrain lymphocyte exhaustion while mitigating CV irAEs consequent to PD-1 blockade.

## Introduction

Immune checkpoint inhibitors (ICI)s targeting programmed cell death protein 1 (PD-1), its ligand PDL-1, or cytotoxic T-lymphocyte antigen 4 (CTLA-4) reinvigorate exhausted CD8^+^ T cell responses against tumor cells (1). ICIs have shown remarkable clinical benefit in a variety of tumors however many cancer patients do not respond or relapse after ICI monotherapy (2). Therapeutic combinations to potentiate and broaden the clinical responses to ICIs are already in use or under clinical investigation (3).

Prostaglandin (PG) E**_2_**, generated by the sequential action of the cyclooxygenase (COX) and microsomal PGE synthase (mPGES-1) enzymes, has immunosuppressive effects in the tumor microenvironment (TME), acting through cognate E2 (EP2r) or 4 receptors (EP4r) expressed on immune cells (4, 5). Specifically, EP2r and EP4r are co-expressed with PD-1 and CLTA-4 on exhausted CD8^+^ T-cells (6) and their activation suppress CD8^+^ T-cell survival and cytotoxicity(7). Prima and colleagues (8) have reported that tumor-associated macrophages (TAMs) expressing high PD-L1 produced more tumor suppressive PGE**_2_**, concomitant with elevated expression of Cox-2 and mPges-1. Pharmacological inhibition of COX-2 reduced PGE**_2_** production by tumor cells *in vitro* and suppressed PD-L1 expression by orthoptic tumor tissues in mice in vivo.

This has rationalized the use of drugs that block PGE_2_ biosynthesis or function in combination with ICIs. Clinical trials are currently underway to explore the potential advantages of combining ICIs with COX inhibitors (NCT03026140, NCT03926338, NCT03728179) or EPr4 antagonists (NCT05944237, NCT05205330) in cancer. However, it is unclear how such combinations might the influence the incidence of CV irAEs in cancer patients, such as myocarditis, atherosclerosis, and thrombosis(9). As the ongoing clinical trials are inadequately powered to address this question, we have focused on one such CV irAE, atherosclerosis, to address this question. Depletion of Pd-1 or Pd-L1/L2 accelerates the progression of atherosclerosis in hyperlipidemic mice, coincident with the expansion of the macrophage and T cell content of aortic plaques (10–12). Conversely, overexpression of CTLA-4 or activation of PD-1 pharmacologically restrains atherogenesis and immune cell infiltration into the plaques (13, 14). We have previously demonstrated that depletion of mPges-1 globally or specifically in myeloid cells restrains atherogenesis in hyperlipidemic mice (Ldlr^-/-^). Similarly, Ep4r deletion in hematopoietic cells delays early atherogenesis in Ldlr^-/-^ mice (15). Here, we sought to study the impact of varied approaches to disruption of the PGE_2_ synthesis / response pathway on the accelerated atherogenesis that occurs in hyperlipidemic mice consequent to Pd-1 depletion.

## Materials and methods

All reagents used were purchased from MilliporeSigma (St. Louis, MO) unless otherwise stated. A detailed description of the experimental methods is provided in the Supplemental Methods.

### Generation of mPges-1 and EPr4 -deficient mice

All mice used were on a C57BL/6J background. The Pd-1^-/-^ mouse line was kindly provided by Dr. John E. Wherry from the University of Pennsylvania (16, 17). The EPr4^F/F^ mouse line was a kind gift of Dr. Richard M. Breyer from the Vanderbilt University Medical Center (18).

All the mice used in the current study were on a Ldlr^-/-^ / C57BL/6J background, purchased from the Jackson Laboratory unless otherwise stated. Male Pd-1^-/-^ mice were mated with female Ldlr^-/-^ mice to obtain Pd-1^-/-^/ Ldlr^-/-^ mice for breeding with mPges-1^-/-^, Lysozyme M (LysM)-Cre^+/-^, mPges-1^F/F^, CAGGCre-ERTM (Ind-Cre^+/-^), and Ep4r^F/F^ mice. Briefly, male mPges-1^-/-^ mice were mated with female Pd-1^-/-^/ Ldlr^-/-^ mice to select mPges-1^+/+^ Pd-1^-/-^/ Ldlr ^-/-^ as controls and mPges-1^-/-^/ Pd-1^-/-^/ Ldlr ^-/-^ as global mPges-1^-/-^ mice. The same breeding strategies were followed to obtain myeloid-specific mPges-1-deficient mice using LysM-Cre^+/-^ and mPges-1^F/F^ mice (19) on a Pd-1^-/-^/ Ldlr^-/-^ background. We used LysM-Cre^+/-^/ mPges-1^+/+^/ Pd-1^-/-^/ Ldlr^-/-^ as controls and LysM-Cre^+/-^/ mPges-1^F/F^/ Pd-1^-/-^/ Ldlr^-/-^ as myeloid-specific mPges-1^-/-^ mice. For global Epr4 deficient mice on Pd-1^-/-^/ Ldlr^-/-^ backgrounds, male Ind-Cre**^+/-^** mice were mated with female Epr4^F/F^ mice to generate Ind-Cre**^+/-^**/ Epr4^F^**^/+^** mice. After several cycles of breeding, male Ind-Cre**^+/-^**/ Epr4^F^**^/+^**/ Pd-1^-/-^/ Ldlr^-/-^ mouse lines were crossed with female Epr4^F/+^/ Pd-1^-/-^/ Ldlr^-/-^ to generate Ind-Cre**^+/-^**/ Pd-1^-/-^/ Ldlr^-/-^ or Epr4^F/F^/ Pd-1^-/-^/Ldlr^-/-^ (as control mice) and Ind-Cre**^+/-^**/ Epr4^F^**^/^**^F^/ Pd-1^-/-^/ Ldlr^-/-^ (as global Epr4^-/-^ mice). Tamoxifen dissolved in corn oil (100mg/ Kg BW, oral gavage, 5 days) was used to induce Epr4 deletion. The presence of mPges-1, Pd-1, LysM-Cre^+/-^, Ind-Cre^+/-^, flox flanking mPges-1 and Epr4, and Ldlr alleles were assessed by polymerase chain reaction (PCR) analyses. Their genetic background was characterized by a 128-SNPs panel composed of loci chosen to differentiate C57Bl/6 sub-strains. This revealed that our mouse models were between 45-69 % of C57Bl/6J background (mPges-1^-/-^/ Pd-1^-/-^/ Ldlr^-/-^: 60.1%; LysM-Cre^+/-^/ mPges-1^FF^/ Pd-1^-/-^/ Ldlr^-/-^: 45.2%; Ind-Cre^+/-^/ Epr4^FF^/ Pd-1^-/-^/ Ldlr^-/-^: 69.2%). All animal housed and the procedures were conducted in accordance with National Institutes of Health regulations and were approved by the Institutional Animal Care and Use Committee (protocol #804754) of the University of Pennsylvania. Animals that developed skin lesions during the feeding of a high fat diet were excluded.

### Statistics

The residuals were tested for normality by the D’Agostino/ Pearson or Shapiro-Wilk test. Post-hoc analysis was performed by pairwise t-tests, with the Bonferroni correction unless otherwise stated. All significant tests were further validated by non-parametric Mann-Whitney and Wilcoxon tests to make sure they were not biased by parametric assumptions. A significance threshold of 0.05 was used for all tests after correcting for multiple testing. Sample sizes (10–20) were based on power analysis of the test measurement and the desire to detect a minimal 10% difference in the variables assessed with α = 0.05 and the power (1-β) = 0.8.

## Results

### Deletion of mPges-1 globally or specifically in myeloid cells restrains atherogenesis augmented in Pd-1^-/-^/ Ldlr^-/-^ mice fed a Western Diet

As expected, deletion of Pd-1 accelerated the accumulation of atherosclerotic plaques in male and female Ldlr^-/-^ mice fed a WD for 24 and 36 weeks, respectively (Figure 1A). This augmented atherosclerotic phenotype was restrained by deletion of mPges-1 globally (Figure 1B), and specifically in myeloid cells (LysM-Cre^+/-^, Figure 1C) in both sexes.

**Figure 1.**
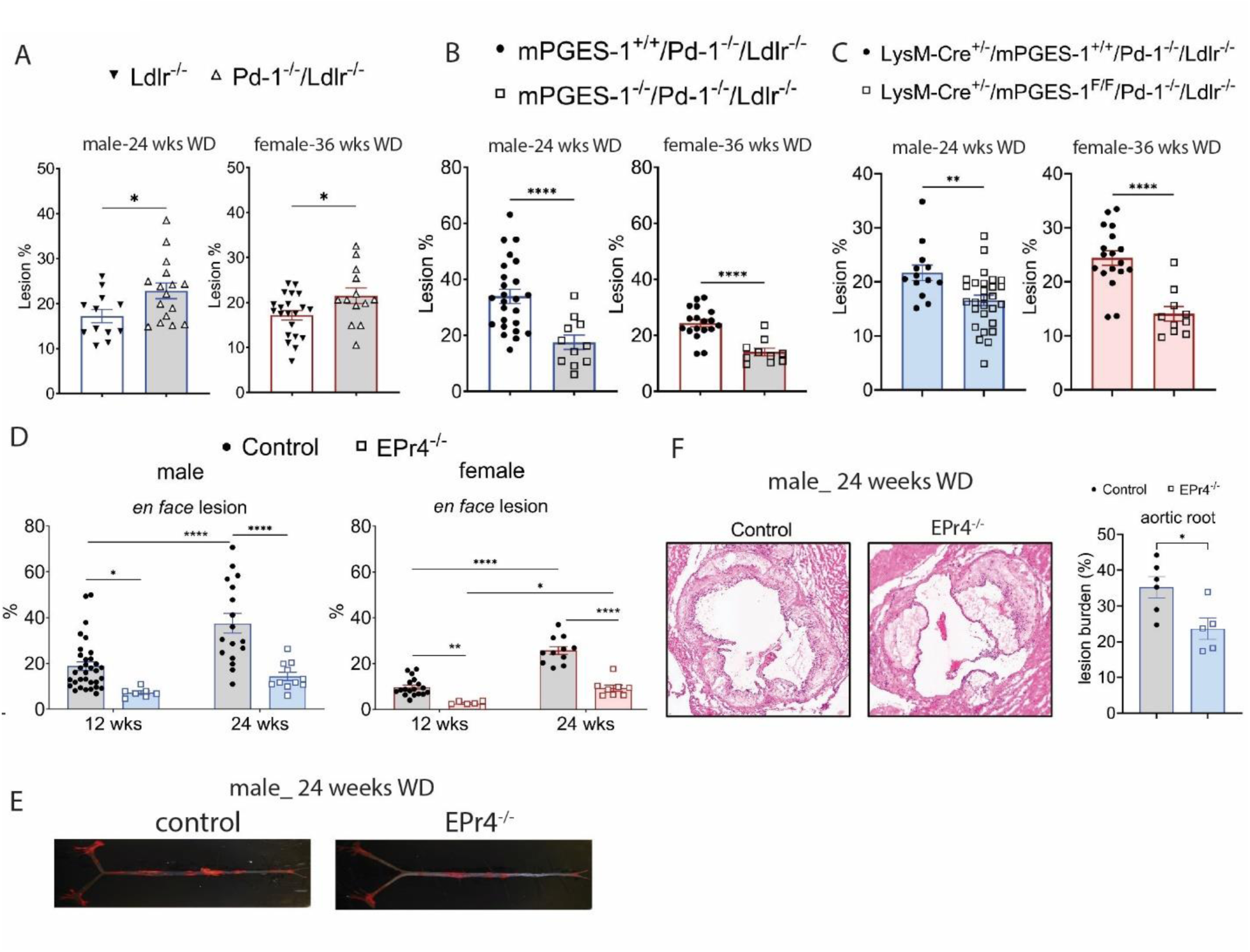
Depletion of mPges-1, globally and in myeloid cells, and EPr4 restrain atherogenesis in Pd-1^-/-^/ Ldlr^-/-^ mice fed a WD. Aortic atherosclerotic lesion burden, represented by the percentage of lesion area to total aortic area, was quantified by *en face* analysis of aortas stained with Sudan IV from mice fed a WD. (A) Depletion of Pd-1 accelerated atherogenesis in Ldlr^-/-^ mice fed a WD for 24 (male) and 36 weeks (female). Depletion of mPges-1 globally (B), specifically in myeloid cells (C), and EPr4 globally (D) restrained the accumulation of atherosclerotic plaques in Pd-1^-/-^ /Ldlr^-/-^ mice of both sexes. (F) Aortic root lesion burden was quantified by H&E staining of cross-sections of OCT embedded aortas from male Pd-1^-/-^/Ldlr^-/-^ mice on a WD diet for 24 weeks. Representative cross-sectional images are shown (right panels). Depletion of EPr4 significantly reduced aortic root lesion burdens. (E) Images of Sudan IV stained e*n face* aortas of control and EPr4^-/-^ male mice on a WD diet for 24 weeks. For A, B, C and D, a Mann Whitney test (2-tailed) revealed a significant effect of genotypes (Pd-1 or mPges-1) on aortic *en face* lesion burden, and EPr4 on aortic root lesion burden. Data are expressed as means ± SEMs. \**p*< 0.05, \*\**p*< 0.01, \*\*\*\**p*< 0.0001, n= 5-6 per group. For D, two-way ANOVA revealed that aortic lesion burden was significantly affected by feeding time or deletion of EPr4. Bonferroni multiple comparison test was used to test for significant differences between controls and EPr4^-/-^ mice and feeding time. Data are expressed as means ± SEMs. \**p*< 0.05, \*\**p*< 0.01, \*\*\*\**p*< 0.0001; n=6-31 per group. Control-Ind-Cre^+/-^/Pd-1^-/-^/Ldlr^-/-^ or EPr4^F/F^/ Pd-1^-/-^/Ldlr^-/-^, EPr4^-/-^-Ind-Cre^+/-^/ EPr4^F/F^/Pd-1^-/-^/Ldlr^-/-^.

Both male and female mPges-1^-/-^ mice gained significantly less weight compared to their littermate controls fed a WD (Figure S1A). Except for heart rate (HR) in mPges-1^-/-^ female mice, systolic blood pressure (SBP) and HR were not significantly altered (Figure S1B-S1C). Plasma levels of total cholesterol, non-high-density lipoprotein cholesterol (non-HDL-C) and glucose (Figure S2A-S2E) were lower in these mice than in controls. This was also true of mice lacking Epr4 (Figures S3A-3C, S4A and S4B) but while depletion of Epr4 reduced lipoprotein lipase activity (LPL; Figure S3D) in female mice fed a WD, this was unaltered by deletion of mPges-1 globally in male mice fed a WD for 12 (Figure S3E).

Deletion of mPges-1 in myeloid cells (LysM-Cre^+/-^) did not significantly alter weight gain, SBP, HR, plasma total cholesterol, non-HDL-C, HDL-C, triglycerides (TGs), or glucose levels after feeding a WD (Figures S1D-F and S2F-J). The results in the global mPges-1^-/-^ mice are consistent with the findings as reported by Ballesteros-Martίnez et al. (20).

### Depletion of Epr4 restrains atherosclerosis-prone Pd-1^-/-^/ Ldlr^-/-^ mice fed a WD

From here onwards, we refer to Ind-Cre^+/-^/ Pd-1^-/-^/ Ldlr^-/-^ or Epr4^F/F^**/** Pd-1^-/-^/ Ldlr^-/-^ mice as controls and Ind-Cre^+/-^/ Epr4^F/F^**/** Pd-1^-/-^/ Ldlr^-/-^ as EPr4^-/-^. The effect of tamoxifen-induced recombination of EPr4 depletion on the expression of Epr1, -r2, -r3, -r4, Cox-1, Cox-2, and mPges-1 in multiple tissues is displayed on a heatmap (Figure S5).

Depletion of Epr4 significantly restrained the progression of atherosclerosis in mice of both sexes at 12 and 24 weeks of feeding a WD, as reflected by *en face* analyses of aortic plaques stained with Sudan IV (Figure 1D). Representative *en face* whole aortic lesion images stained with Sudan IV dye of male mice on 24 weeks of a WD are shown (Figure 1E).

Consistent with the *en face* analyses, aortic root lesion burden, as reflected by H&E staining, was significantly decreased in Epr4^-/-^ male mice at 24 weeks on a WD (Figure 1F).

Depletion of Epr4 did not significantly alter body weight, SBP and HR of mice of both sexes on 12 and 24 weeks of feeding a WD (Table S1).

### Differential impact of mPges-1 and Epr4 depletion of prostanoid biosynthesis in Pd-1^-/-^/ Ldlr^-/-^ mice

As expected, deletion of mPges-1 significantly reduced PGE_2_ biosynthesis in male mice and trended to reduced PGE_2_ biosynthesis in female mice, as reflected by urinary PGE_2_ metabolite excretion (PGEM, Figure 2A-2B). Urinary PGIM, PGDM and TxM were not significantly altered. However, urinary PGIM was significantly elevated in mice deficient of mPges-1 in myeloid cells, (Figure 2C-2D). Urinary PGEM, PGDM and TxM were not significantly changed between controls and myeloid mPges-1 -deficient mice after feeding a WD, likely reflecting the minor contribution of the myeloid compartment to the systemic biosynthesis of prostanoids.

**Figure 2.**
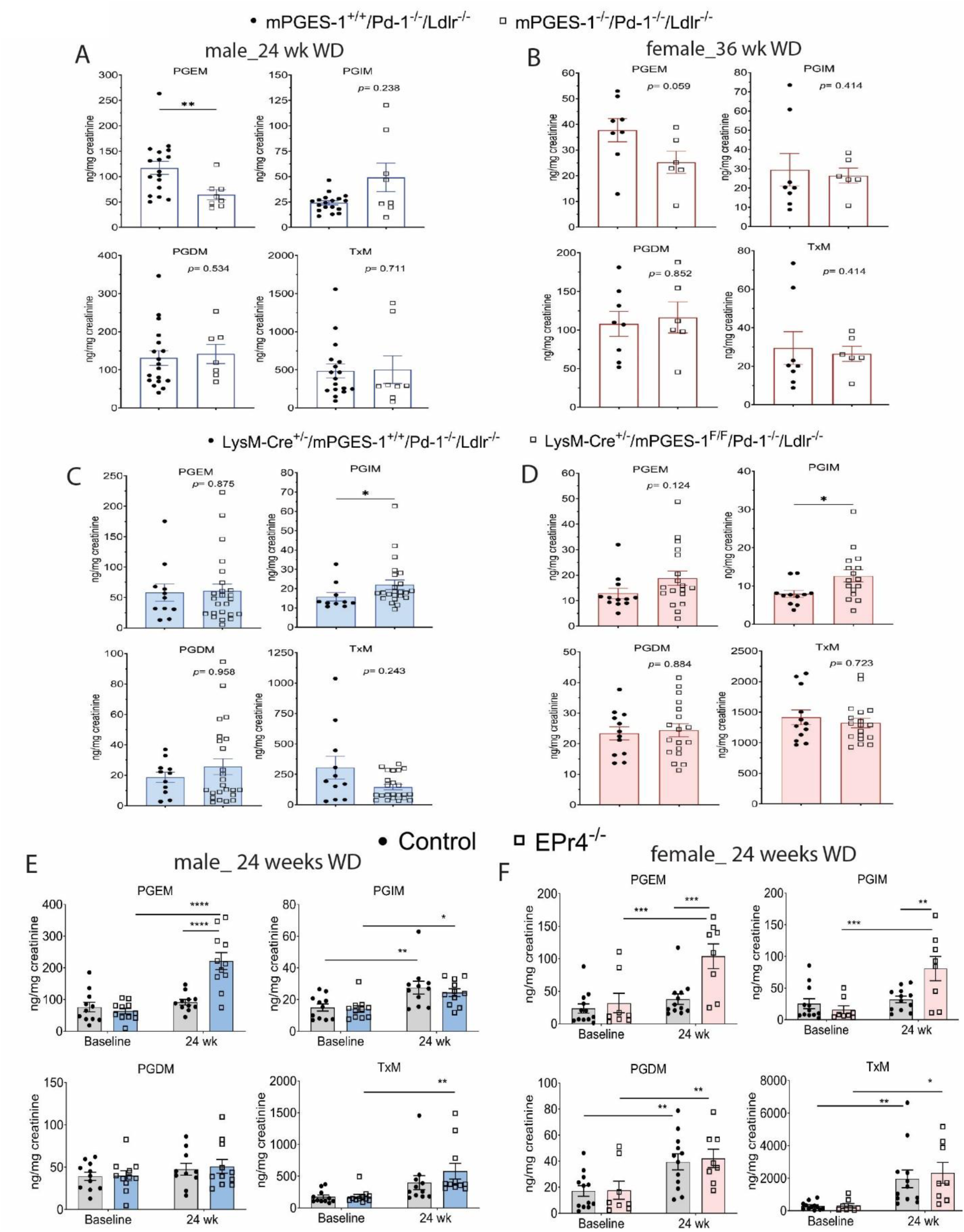
Disruption of PGE_2_/EPr4 response pathways significantly alter urinary prostanoid metabolites in Pd-1^-/-^/Ldlr^-/-^ mice fed a WD. Depletion of mPges-1 significantly reduced PGE_2_ biosynthesis in Pd-1^-/-^/Ldlr^-/-^ mice whilst EPr4 depletion significantly elevated PGE_2_ and PGI_2_ biosynthesis in female Pd-1^-/-^/Ldlr^-/-^ mice fed a WD. Fasting (10am-5pm) urine samples from mice were collected before and after feeding mice a WD, and prostanoid metabolites (PGEM-7-hydroxy-5, 11-diketotetranorprostane-1, 16-dioic acid, PGIM-2, 3-dinor 6-keto PGF_1α_, TxM-2, 3-dinor TxB_2_, tetranor PGDM-11, 15-dioxo-9_α_-hydroxy-2, 3, 4, 5-tetranorprostan-1, 20-dioic acid, M-metabolite) were analyzed by LC/MS/MS as described in the Supplemental Methods. (A and B) Depletion of mPges-1 significantly reduced PGE_2_ biosynthesis in Pd-1^-/-^/Ldlr^-/-^ mice fed a WD for 24 weeks (male) and female (36 weeks). (C and D) Depletion of mPges-1 in myeloid cells significantly increased PGI_2_ biosynthesis in Pd-1^-/-^/Ldlr^-/-^ mice fed a WD for 24 weeks (male) and female (36 weeks). (E and F) Depletion of EPr4 significantly increased PGEM and PGIM in female mice after feeding a WD. In male mice, only PGEM was significantly increased in EP4r^-/-^ mice after feeding a WD. For A, B, C and D, a Mann Whitney test (2-tailed) revealed a significant effect of genotype on urinary PGM. Data are expressed as means ± SEMs. \**p*< 0.05, \*\**p*< 0.01. For E and F, 2-way ANOVA showed that urinary prostanoid metabolites were significantly affected by feeding a WD or EPr4 globally. Bonferroni multiple comparison test was used to test for significant differences between controls and EPr4^-/-^ mice. Data are expressed as means ± SEMs. \**p*< 0.05, \*\**p*< 0.01, \*\*\**p*<0.001, n= 6-24 per genotype. Control-Ind-Cre^+/-^/Pd-1^-/-^/Ldlr^-/-^ or EPr4^F/F^/ Pd-1^-/-^/Ldlr^-/-^, EPr4^-/-^-Ind-Cre^+/-^/ EPr4^F/F^/Pd-1^-/-^/Ldlr^-/-^.

In contrast to mPges-1-deficient mice, depletion of Epr4 significantly augmented both urinary PGEM and PGIM in female mice after feeding a WD for 24 weeks (Figure 2E). Both urinary PGDM and TxM increased on a WD but were unaltered by genotype (Figure 2F).

The increase in PGE_2_ biosynthesis in the face of Epr4 depletion likely represents a compensatory augmentation of agonist production driven by increased expression of Ptgs-1 and mPges-1 genes and PTGS-1 and mPGES-1 protein in aorta (Figure 3A-3C) of Epr4^-/-^ mice. Consistent with these results, inhibition of PTGS-1 with SC-560 (25) but not of PTGS-2 with nimesulide significantly reduced urinary PGEM in EPr4^-/-^ mice compared to controls on a WD (Figure S6A and 6B). SC560 also abrogated but did not abolish the difference in WBCs and lymphocytes by genotype in mice fed a WD (Figure S6C and S6D).

**Figure 3.**
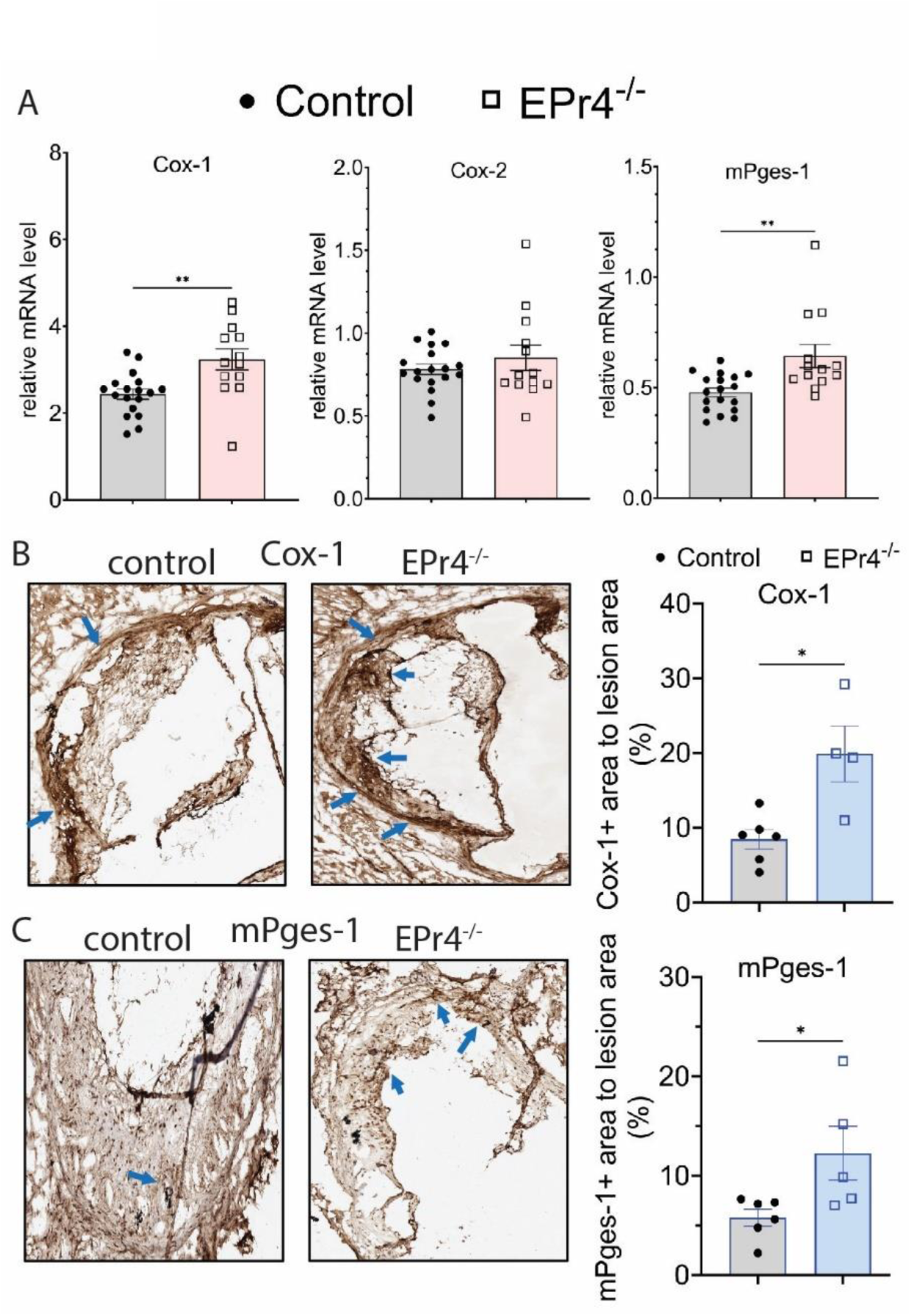
Depletion of Epr4 increases aortic Cox-1 and mPges-1 mRNA and protein levels in mice fed WD for 12 weeks. Female mice fed a WD for 12 weeks were used to collect whole aorta for RNA extraction as detailed in the Supplemental Methods. Cox-1, Cox-2 and mPges-1 mRNA levels of aortas were measured by rt-qPCR. (A) Depletion of Epr4^-/-^ increased Cox-1 and mPges-1 mRNA levels in mouse aorta from female mice fed a WD for 12 weeks. Data are expressed as means ± SEMs. \*\**p*< 0.01, n= 13-18 per group. (B-C) Immunohistochemical analysis of aortic root lesion displayed a significant increase in Cox-1 or mPges-1 positive staining in EPr4 deficient aortic root lesions. Representative cross-sectional images of aortic root lesions are shown. Arrows indicate Cox-1 or mPges-1 positive area. Cox-1 or mPges-1 positive area was analyzed using Image-Pro Plus software. \**p*< 0.05, n= 4-6 per group. A Mann Whitney test (2-tailed) revealed a significant effect of Epr4 deletion on Cox-1 and mPges-1 transcripts and protein expression levels. Data are expressed as means ± SEMs. Control-Ind-Cre^+/-^/Pd-1^-/-^/Ldlr^-/-^ or EPr4^F/F^/ Pd-1^-/-^/Ldlr^-/-^, EPr4^-/-^-Ind-Cre^+/-^/ EPr4^F/F^/Pd-1^-/-^/Ldlr^-/-^.

### Differential impact of mPges-1 and Epr4 deletion on the immune-cell infiltration in the atherosclerotic plaques

Previous studies have shown that deletion of Pd-1 or Pd-L1/L2 accelerated atherogenesis in Ldlr^-/-^ mice coincident with the accumulation of macrophages, CD4^+^ and CD8^+^ T cells in aortic plaques (11, 12). To investigate the impact of mPGES-1 deletion on the immune cells present in the aortic plaques, we performed flow-cytometry analysis on CD45^+^ cells isolated from the aortas of male Pd-1^-/-^/Ldlr^-/-^ mice fed a WD for 12 weeks (Figure 4A). mPGES-1 deletion did not alter the infiltration of CD8^+^ T cells and macrophages into the aorta of Pd-1^-/-^/Ldlr^-/-^ mice but reduced the infiltration of neutrophils (Figure 4A-4B).

**Figure 4.**
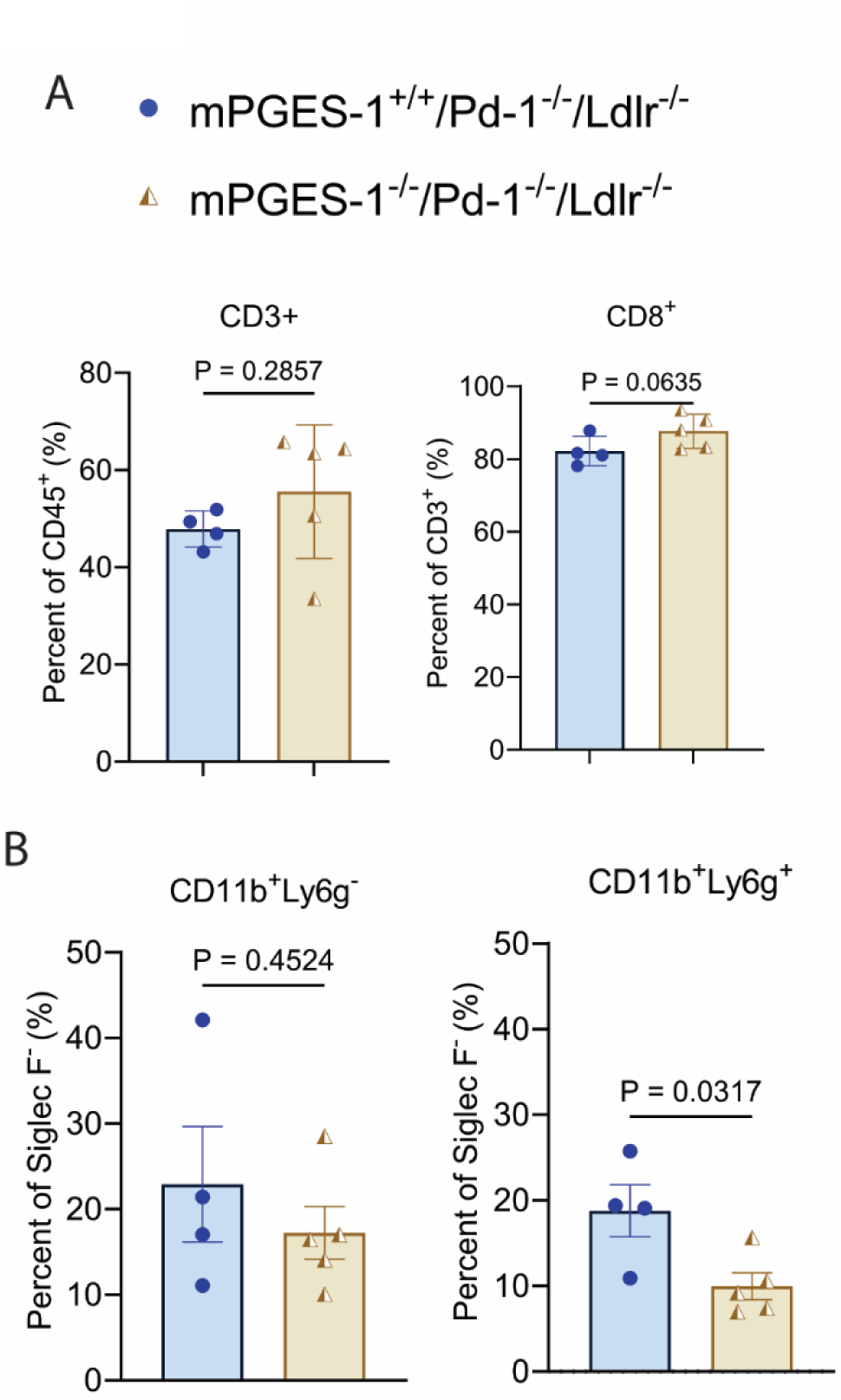
Deletion of mPges-1 reduces the neutrophil population in aortic plaques of male mice fed a WD for 12 weeks. Immune cells in atherosclerotic plaques of mice fed a WD was isolated from digested aortas. Staining of immune cells with antibodies was described in the Supplemental method. (A) Deletion of mPges-1 did not significantly change the CD8^+^ T cell population in the aortic plaques. (B) Deletion of mPges-1 reduced neutrophils (CD11b^+^Ly6g^+^) in the aortic plaques. Mann Whitney test (2-tailed) revealed a significant effect of genotype on neutrophil population in the aortic plaques. Data are expressed as means ± SEMs. \**p*< 0.05, n= 4-5 per genotype.

To investigate further the cellular immune landscape of the aortic plaques, we performed sc-RNA sequencing, using CD45^+^ immune cells from atheroprone aortas of female mice fed a WD for 12 weeks (Figure 5A).

**Figure 5.**
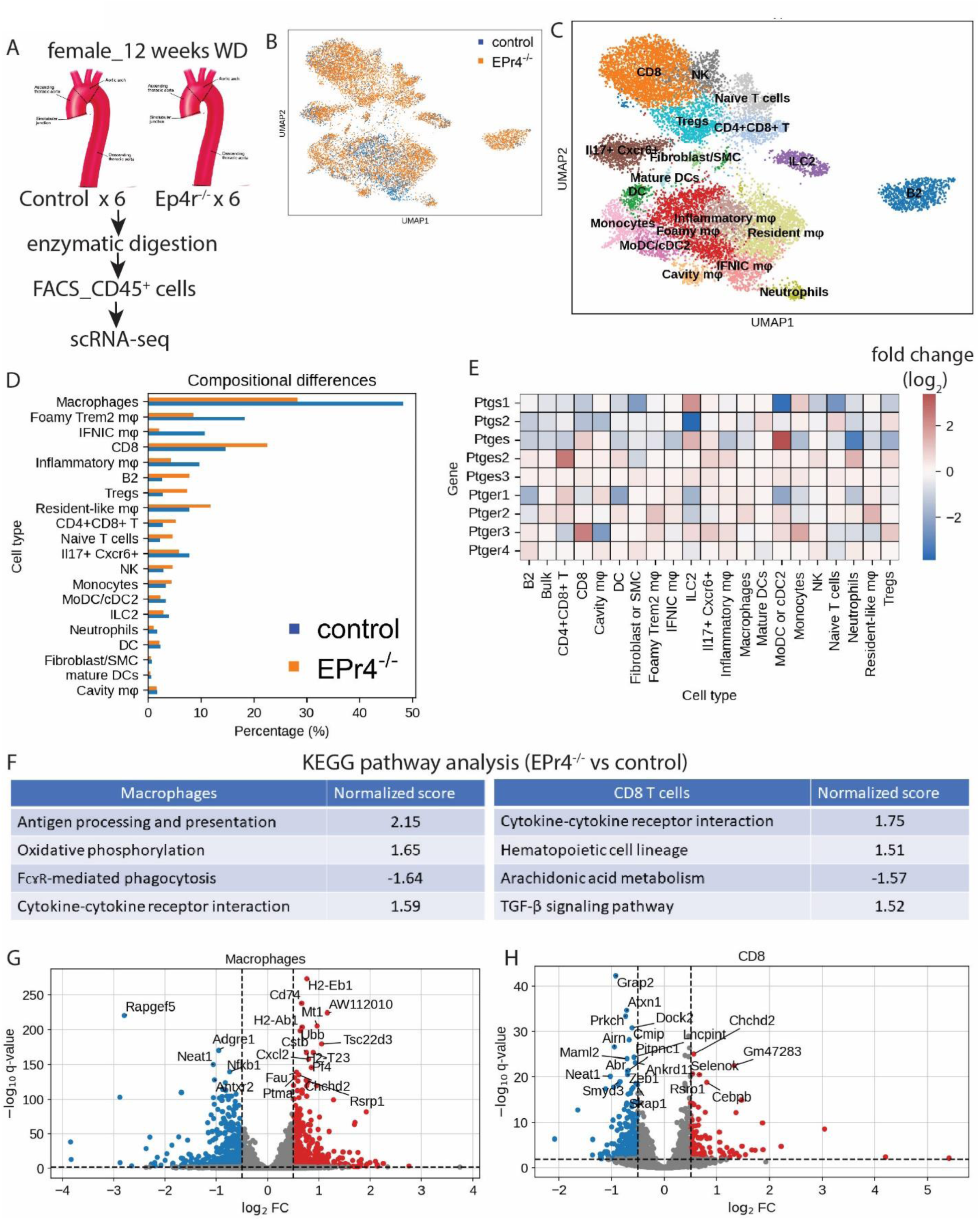
Single-cell RNAseq analysis identified 19 clusters of CD45^+^ cells and revealed a reduction in macrophages and an increase in T cells in Ep4r^-/-^ cells from mouse atherosclerotic lesions. (A) Experimental design of scRNA sequencing of CD45^+^ cells in atherosclerotic plaques of female mice fed a WD for 12 weeks. Digested cells from six atherosclerotic aortas were pooled to yield enough cells for scRNA-sequencing. (B) UMAP representation of aligned gene expression data in single CD45^+^ cells isolated from atherosclerotic plaques of controls (blue) and EPr4^-/-^ (orange) female mice fed a WD for 12 weeks. (C) UMAP showing the distribution and identify of the nineteen different cell clusters from mouse atherosclerotic lesions. CD8^+^ T cells, NK cells, naïve T cells, Treg cells, CD4^+^CD8^+^ T cells, IL17^+^Cxcr6^+^, fibroblast/SMC, ILC2, mature DC, DC, B2, monocytes, inflammatory M_ϴ_, foamy Trem2 M_ϴ_, resident-like M_ϴ_, IFNIN M_ϴ_, cavity M_ϴ_, neutrophils. (D) Compositional differences of defined cell populations in each cluster as defined in C. Macrophages populations decreased and CD8 T cells increased in EPr4^-/-^ mice compared to controls mice. (E) Heatmap showing differential gene expression of prostanoid pathways. (F) KEGG pathway analysis of differentially expressed genes in macrophages (all five clusters of macrophages) and CD8^+^ T cells. Positive value represents upregulation of pathways, negative value represents downregulation of pathways. Volcano plot of the top differentially expressed genes in all macrophages (G) and CD8^+^ T cells (H). Control-Ind-Cre^+/-^/Pd-1^-/-^/Ldlr^-/-^ or EPr4^F/F^/ Pd-1^-/-^/Ldlr^-/-^, EPr4^-/-^-Ind-Cre^+/-^/ EPr4^F/F^/Pd-1^-/-^/Ldlr^-/-^.

Following the annotations as reported by Zernecke et al. (21) and Zhang et al. (22), scRNA-seq analyses revealed 19 distinct cell clusters with unique marker genes as displayed in UMAPs (Figure 5B-CD45^+^ atherosclerotic cells from control and EPr4^-/-^ mice, Figure 5C- 19 cell clusters, Figure S7A- heatmap of marker genes of each cell cluster). As shown in Figure 5C, macrophages and T cells were the dominant immune cells isolated. Five clusters of macrophages (Mφ); resident-like Mφ, inflammatory Mφ, foamy triggering receptor expressed on myeloid cell 2 (foamy Trem2) Mφ, interferon inducible (IFNIC) Mφ and cavity Mφ were identified. Similarly, we found five clusters of T cells, namely, CD8^+^ T cells, CD4^+^CD8^+^ T cells, Il17^+^Cxcr6^+^ T cells, naïve T cells and Treg cells. Other identified cell clusters including B2 cells, neutrophils, monocytes, monocyte-derived dendritic cells/ conventional dendritic cells type 2 (MoDC/ cDC2), plasmacytoid DCs (DCs), mature DCs, natural killer (NK) cells, type 2 innate lymphocyte cells (ILC2) and fibroblast/ smooth muscle cells (SMCs).

Depletion of Epr4 restrained the accumulation of foamy Trem2 Mφ, IFNIC Mφ, inflammatory Mφ and cavity Mφ, but not resident-like Mφ, in aortic plaques as compared to control mice during the development of atherosclerosis (Figure 5D). Conversely, four of the five T cell clusters increased in proportions in the aortic plaques of Epr4^-/-^ mice, as well as for B2 cells (Figure 5D). Given that foamy Trem2, IFNIC and inflammatory Mφ exhibit overlapping characteristics, they expressed varying levels of Ccl2, Ccl4, Ctsd, Lgals3 and Wfdc17 (Figure S7A). Depletion of Epr4 altered expression of Ptgs1 (Cox-1), Ptgs2 (Cox-2), Ptges (mPges-1), Ptges2 (mPges-1), Ptges3 (cPges), and the receptors of PGE_2_ in all cell types to different degrees (Figure 5E). For example, Ptges2 (mPges-2), Ptger3 (EPr3), Ptgs1 (Cox-1) and Ptges (mPges-1) were upregulated in CD4^+^CD8^+^ T cells, CD8^+^ T cells, IL-C2 and MoDC/cDC2 cells, respectively. As expected, Epr4 gene expression was reduced across the different cell types.

KEGG pathway analyses associated with changes in gene expression of macrophages (all five types) showed that antigen processing and presentation, oxidative phosphorylation, FcɤR- mediated phagocytosis, and cytokine-cytokine receptor interaction were differentially altered between cells from control and Epr4^-/-^ mice (Figure 5F, left table). For CD8^+^ T cells, pathway enrichment analyses revealed that cytokine-cytokine receptor interaction, hematopoietic cell lineage, arachidonic acid metabolism and TGF-β signaling pathway were differentially expressed and modulated by Epr4 depletion (Figure 5F, right table). As shown in Figure 5G and 5H, depletion of Epr4 resulted in more differently expressed genes (DEGs) in macrophages than in CD8^+^ T cells as displayed in Volcano plots. In macrophages, increased expression of AW112010, a long noncoding RNA (IncRNA), promoted the expression of major histocompatibility complex class I/ II (MHC I/ II) molecules including H2-EB1, H2-Ab1, and H2-T23, which are involved in mitigating immune responses and presenting antigen to immune cells including T cells. Another lncRNA, nuclear paraspeckle assembly transcript 1 (NEAT1) was reduced in EPr4^-/-^ macrophages and CD8^+^ T cells, coincident with the restraint of atherosclerotic plaques. Similar findings of DEGs were observed in the subtypes of macrophages and T cells (Figure S7B-J).

### Depletion of Epr4 impairs chemotaxis of bone marrow derived macrophages (BMDMs) and restrains the accumulation of aortic macrophages

To address further the reduction in macrophages in aortic plaques of Epr4^-/-^ mice, we studied their bone marrow derived macrophages (BMDMs). BMDMs from female Epr4^-/-^ mice displayed significantly impaired chemotaxis as compared to control BMDMs (Figures 6A and B) irrespective of being fed a standard laboratory diet (SLD) or a WD for 12 weeks while cell viability was preserved (Figure 6C).

**Figure 6.**
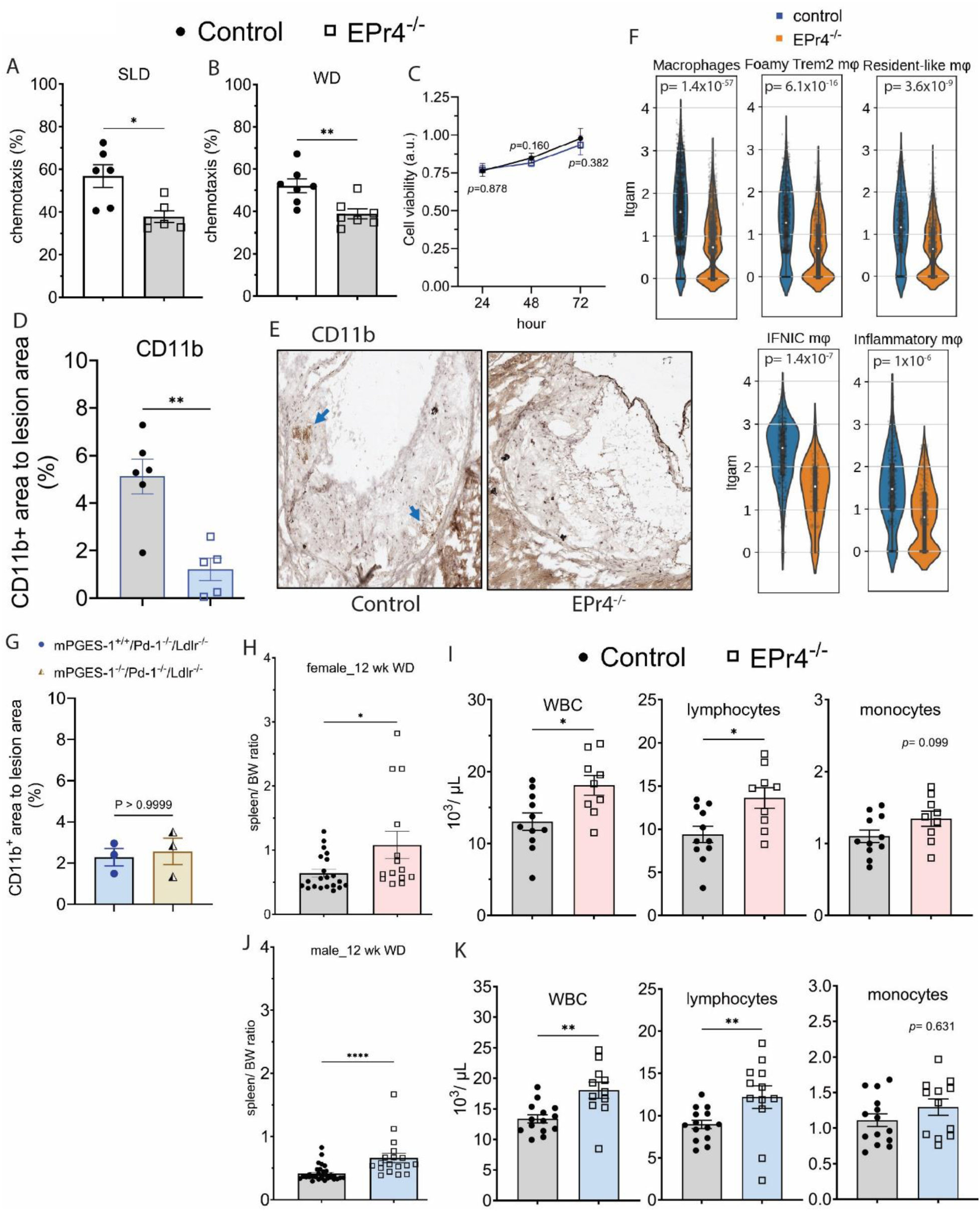
Depletion of EPr4 impairs chemotaxis of bone-marrow derived macrophages (BMDMs), reduces aortic macrophage content, and stimulates extramedullary hematopoiesis. Bone-marrow derived macrophages (BMDMs) were isolated from control and EPr4 deficient mice fed a standard laboratory diet (SLD) or a WD for 12 weeks, to perform chemotaxis assay using Boyden chambers. (A and B) Depletion of EPr4 significantly impaired chemotaxis of BMDMs in Boyden chambers for 48 hours. The presence of a cocktail of chemotactic factors (LPS- 0.25 µg/mL, oxLDL- 5µg/mL, MCP-1- 50 pg/ML, MIP-1α- 1.5 pg/ML, platelet-derived growth factor (PDGF)- 5 ng/mL) did not confer additional migration of BMDMs. Data are expressed as means ± SEMs from two independent experiment, \**p*< 0.05, \*\**p*< 0.01, n= 6- 7 per group. (C) Cell viability was not significantly altered between control and EPr4^-/-^ BMDMs for up to 72 hours. Data are expressed as means ± SEMs, *p*> 0.05, results are representative of two independent experiment, n= 8 replicate per group. (D and E) Immunohistochemical analysis of aortic root lesion displayed a significant decrease in CD11b positive staining, a marker of macrophages, in EPr4 deficient aortic root lesions. Representative cross-sectional images of aortic root lesions are shown. Arrows indicate CD11b positive area. CD11b positive area was analyzed using Image-Pro Plus software. \*\**p*< 0.01, n= 5- 6 per group. (F) Itgam (CD11b) gene expression in macrophage clusters. (G) Immunohistochemical analysis of CD11b positive staining in aortic root lesions of mPGES-1^+/+^/Pd-1^-/-^/ Ldlr^-/-^ and mPGES-1^-/-^/Pd-1^-/-^/ Ldlr^-/-^ male mice fed a WD for 24 weeks. *p>* 0.05, n= 3 per group. (H -J) Spleen size was normalized to body weight and complete blood count (CBC) was performed using a Sysmex Hematology Analyzer. Depletion of EPr4 significantly increased spleen size, numbers of white blood cells (WBC) and lymphocytes in blood. A Mann Whitney test (2-tailed) revealed a significant effect of EPr4 deletion on chemotaxis of BMDMs, CD11b positive area, spleen size, numbers of WBCs and lymphocytes in the blood. For chemotaxis assay, data are expressed as means ± SEMs from two independent experiment. \**p*< 0.05, \*\**p*< 0.01, \*\*\*\**p*< 0.0001, n= 14- 33 per group (spleen size), n= 9- 14 per group (CBC). Control- Ind-Cre^+/-^/Pd-1^-/-^/Ldlr^-/-^ or EPr4^F/F^/ Pd-1^-/-^/Ldlr^-/-^, EPr4^-/-^- Ind-Cre^+/-^/ EPr4^F/F^/Pd-1^-/-^/Ldlr^-/-^.

An immunohistochemical marker of macrophage infiltration, CD11b, was reduced in aortic roots of atherosclerotic EPr4^-/-^ mice (Figure 6D- 6E, arrows indicate positive staining) as was its expression in macrophage clusters (Figure 6F). In contrast, the percentage of CD11b^+^ cells in the aortic lesions was unaltered by mPges-1 deletion (Figure 6G).

### Depletion of Epr4 enhances extramedullary lymphoid hematopoiesis in hyperlipidemic mice fed a WD

PGE_2_, acting via Epr4, has been reported to promote the survival and expansion of the hematopoietic niche (23, 24). The increase in several T cell populations in aortic plaques of Epr4^-/-^ mice are consistent with enhanced extramedullary lymphoid hematopoiesis. Spleen size normalized to body weight of the animals was significantly larger in EPr4^-/-^ mice as compared to controls after feeding a WD for 12 weeks (Figure 6H- female, 6J- male) while WBC, lymphocyte and platelet counts were elevated in EPr4 deficient mice of both sexes (Figure 6I & 6K, platelets- Figure S8A- S8B). Monocytes, red blood cells and reticulocytes were not significantly altered by genotype. (Figure 6H, 6J, Figure S8A- S8B).

By contrast depletion of mPges-1 globally did not significantly affect heart and spleen weights, numbers of WBCs, RBCs, lymphocytes or platelets in whole blood (Figure S9)

### Epr2 receptor blockade confers no additional restraint on the accumulation of atherosclerotic plaques in Epr4 deficient Pd-1^-/-^/ Ldlr^-/-^ mice

To address the potential functional redundancy between EPr4 and EPr2 in driving lymphocyte exhaustion, we compared Epr4 depleted mice with and without treatment with an EPr2 antagonist (PF-04418948, 10 mg/Kg BW) (25). This conferred no additional impact on atherogenesis (Figure S10A), and spleen size remained enlarged (Figure S10B). However, the increases in WBCs and lymphocytes in EPr4^-/-^ mice (Figure 5I and 5K) were abrogated by the EPr2 antagonist (Figure S10C). RBC number decreased and reticulocyte count increased while the platelet count remained higher in the EPr4^-/-^ mice (Figure S10C) despite treatment with the EPr2 antagonist.

## Discussion

We have previously reported that depletion of mPges-1 globally or specifically in myeloid cells restrain atherogenesis in Ldlr^-/-^ mice fed a WD rich in cholesterol and fats (19, 26). The role of EPr4 deletion in atherogenesis is less clear. EPr4 deletion in hematopoietic cells delayed early atherogenesis in Ldlr^-/-^ mice on WD, while its deletion in the bone marrow or myeloid cells did not impact the size of atherosclerotic plaques in mice fed a WD or in diabetic mice (27, 28). In this study, we replicated the accelerated immune atherogenic phenotype reported in Ldlr^-/-^ mice lacking Pd-1 fed a WD, characterized by lesional infiltration of monocytes and lymphocytes (14, 15). We found that depletion of mPges-1 globally or specifically in myeloid-cells restrained the progression of atherosclerosis in Pd-1^-/-^ Ldlr^-/-^ mice. As reported earlier (25), the increase in PGI_2_ might have contributed to this atheroprotective phenotype, consistent with the deletion of IPr in restraining atherogenesis in hyperlipidemic mice (29, 30). Global EPr4 deletion also delayed atherogenesis in the same mouse model, although this was accompanied with an increase in PGEM, which reflects receptor blockade, as seen with EPr4 antagonists in humans (31, 32). Although the two strategies of PGE_2_ suppression mitigated the immune atherogenesis due to Pd-1 deletion, they had a different impact on the type of immune cells infiltrating in the atherosclerotic plaques. Flow-cytometry analysis revealed that mPGES-1 deletion reduced the infiltration of neutrophils in the atherosclerotic plaques in Pd-1^-/-^ mice, while it did not affect the infiltration of other immune cells. In contrast, scRNA sequencing analyses revelated that Epr4 deletion increased CD8^+^ T cells and reduced the number of foamy Trem2, IFNIC and inflammatory Mφ infiltrating in the atherosclerotic plaques of Pd-1^-/-^ mice. This is exemplified by the upregulation of genes (AW112010, H2-Eb1, H2-Ab1, H2-T23) responsible for antigen processing and presentation are unique to macrophages. Neat1 is a lncRNA that regulates NLRP3 and activates the inflammasomes; deletion of Neat1 suppressed ox-LDL induced inflammation in macrophages *in vitro* (33). The different impacts of mPGES-1 and EPr4 deletion on the macrophage population was confirmed by IHC with CD11b staining. Consistently, BMDMs from EPr4^-/-^ mice showed impairment in chemotaxis *in vitro.* These findings provide a mechanistic explanation for the restraint on plaque progression observed in Pd-1^-/-^/Ldlr^-/-^ mice lacking Epr4. PGE_2_, acting via EPr4, facilitates monocyte recruitment, differentiation of these monocytes into inflammatory macrophages in atherosclerotic plaques thereby driving progression of atherosclerosis (34, 35). The additional blockade of EPr2 in Epr4^-/-^ mice did not detectably augment atheroprotection from Epr4 depletion. Coincident with their atheroprotective phenotypes, depletion of both mPges-1 and Epr4 reduced total cholesterol and LDL-cholesterol and caused a slight increase in triglyceride levels. mPGES-1^-/-^ mice also reduced glycemia as it was previously reported (26). A reduction in LPL, splenomegaly and lymphocytosis was detected in EPr4^-/^, but not mPGES-1^-/-^ mice.

Overall, atherosclerosis and other CV irAEs of ICIs are rare (∼1% of cases), but extremely serious (mortality up to 50%). The recognition of PGE_2_ as a checkpoint has prompted combinatorial clinical trials of ICIs with drugs targeting the COX-2-2/mPGES-1/EPr4 pathway. However, despite the cardiotoxicity of COX-2 inhibitors (36) these trials are not adequately sized to detect an increase in CV AEs from PGE_2_ disruptive drug combinations, hence the importance of addressing this issue in preclinical models. Here we provide evidence that mPges-1 or Epr4 deletion mitigates the pro-atherogenic phenotype observed in Pd-1/Ldlr deficient mice, but with important distinctions in terms of both prostanoid biosynthesis and immune response (Figure S11). Future studies will include comparison with targeted deletion of Ptgs-2, determination of whether the results with atherogenesis extend to other models of irAEs and whether combinatorial targeting of COX-2, mPGES-1 or EPr4 differentially impact the efficacy of ICIs.

## Acknowledgements

We thank Elizabeth J. Hennessy (University of Pennsylvania) for sharing chemicals for the BMDM chemotaxis assay. This work was supported by a grant from NIH (HL141912). We thank members of the Translational Core Laboratory at Children Hospital of Philadelphia for assisting with CBC. GAF is the McNeil Professor of Translational Medicine and Therapeutics and Senior Advisor to Calico Laboratories. GAF held a Merit Award from the American Heart Association during the performance of this work.

