## Supplemental Figures for "Disruption of the PGE_2_ synthesis / response pathway restrains atherogenesis in programmed cell death-1 (Pd-1) deficient hyperlipidemic mice"

**Short title for running head: mPGES-1/ $PGE_2$/ Ep4r signaling in $Pd\text{-}1^{-/-}$/ $Ldlr^{-/-}$ mice**

[1, 2#]Emanuela Ricciotti, PhD; [1#]Soon Yew Tang, PhD; [1]Antonijo Mrčela, PhD; [1]Ujjalkumar S. Das, PhD; [1]Ronan Lordan, PhD; [1]Robin Joshi, PhD; [1]Soumita Ghosh, PhD; [1]Justin Aoyama, BSc; [1]Ryan McConnell, BSc; [1]Jianing Yang, PhD; [1,3]Gregory R. Grant, PhD; [1,4]*Garret A. FitzGerald, MD

[1]From the Institute for Translational Medicine and Therapeutics, Perelman School of Medicine, [2]the Department of Systems Pharmacology and Translational Therapeutics, the [3]Department of Genetics, and the [4]Department of Medicine Perelman School of Medicine, University of Pennsylvania.

*Address for correspondence: Garret A. FitzGerald, Institute for Translational Medicine and Therapeutics, Perelman School of Medicine, 10-110 Smilow Center for Translational Research, 3400 Civic Center Blvd, Bldg 421, University of Pennsylvania, Philadelphia, PA 19104-5158. Fax: 215-573-9135 Tel: 215-898-1184

### These authors contributed equally.

All data are available by contacting the corresponding author.

Supplemental Materials

Supplemental Table S1. Body weight gain, systolic blood pressure (SBP) and heart rate (HR) of controls and EPr4$^{-/-}$ mice on a Western diet (WD).

| | female | | | | male | | | |
|---|---|---|---|---|---|---|---|---|
| | 12 weeks WD | | 24 weeks WD | | 12 weeks WD | | 24 weeks WD | |
| | control | EPr4$^{-/-}$ | control | EPr4$^{-/-}$ | control | EPr4$^{-/-}$ | control | EPr4$^{-/-}$ |
| Weight gain (g) | 4.80 ± 0.41 | 4.01 ± 0.60 | 8.05 ± 0.83 | 6.54 ± 0.84 | 8.72 ± 0.36 | 7.57 ± 0.70 | 10.8 ± 1.03 | 8.67 ± 1.24 |
| SBP (mmHg) | 91.8 ± 2.40 | 106 ± 4.02 | 99.9 ± 2.33 | 96.0 ± 3.86 | 101 ± 1.91 | 96.3 ± 2.0 | 96.7 ± 2.05 | 98.8 ± 2.61 |
| HR (bpm) | 657 ± 16 | 676 ± 24 | 686 ± 11 | 706 ± 8 | 638 ± 9 | 632 ± 24 | 679 ± 9 | 679 ± 12 |

Control- Ind-Cre$^{+/-}$/Pd-1$^{-/-}$/Ldlr$^{-/-}$ or EPr4$^{F/F}$/ Pd-1$^{-/-}$/Ldlr$^{-/-}$, EPr4$^{-/-}$- Ind-Cre$^{+/-}$/ EPr4$^{F/F}$/Pd-1$^{-/-}$/Ldlr$^{-/-}$.

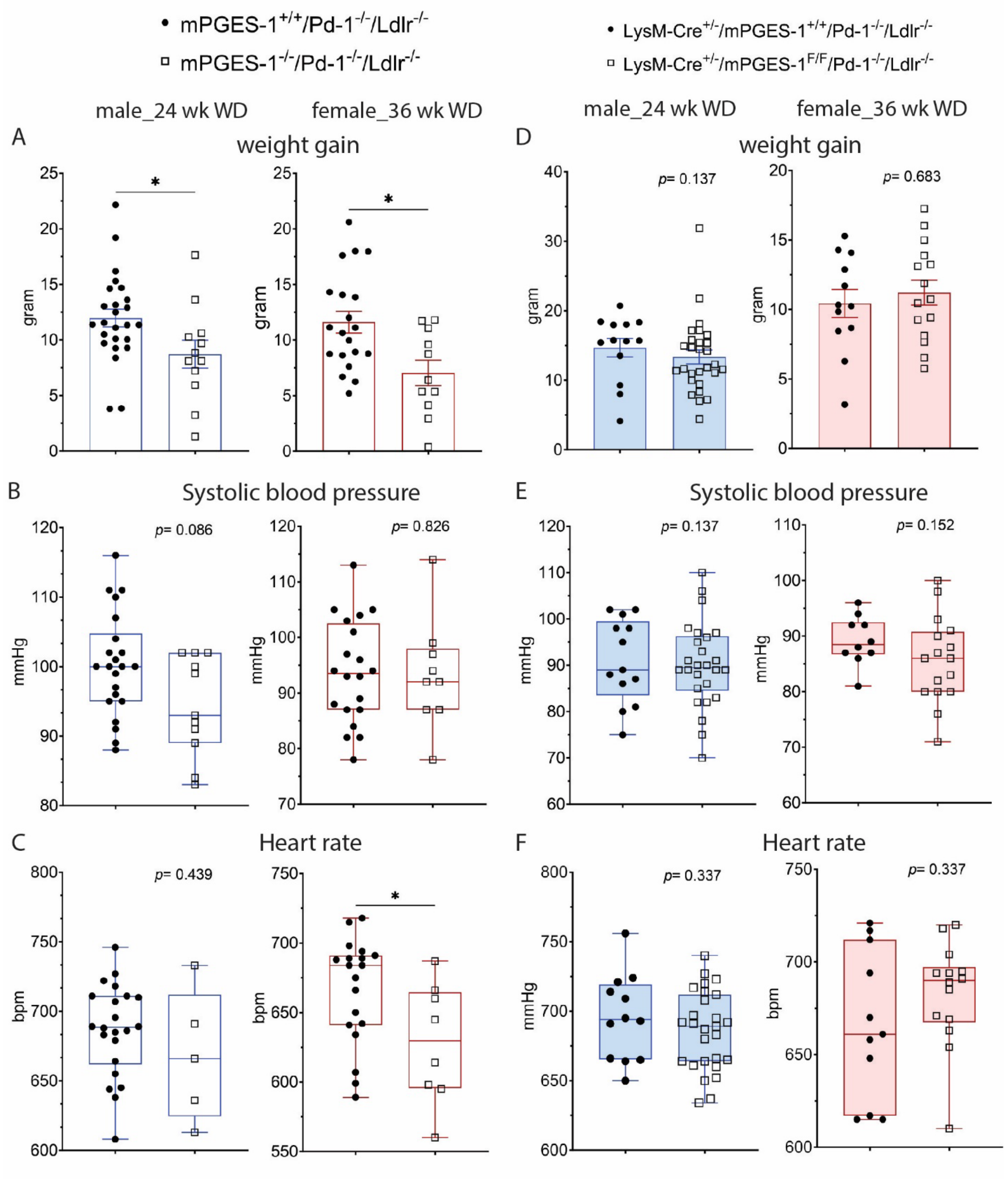


**Supplemental Figure S1. Depletion of mPges-1 globally significantly altered weight gain and heart rate (HR) in Pd-1$^{-/-}$/ Ldlr$^{-/-}$ mice fed a WD.**

(A) Body weight (BW) was calculated based on the difference in BW before and after feeding of a WD. (B) Systolic blood pressure (SBP) and (C) heart rate (HR) were measured using a tail-cuff system in mice fed a WD for 24 weeks (male) and 36 weeks (female). Except female mice, there were no significant differences in SBP and HR between Pd-$1^{-/-}$/$Ldlr^{-/-}$ and mPges-$1^{-/-}$/Pd-$1^{-/-}$/$Ldlr^{-/-}$ mice of both sexes (A- weight gain, B- SBP, C- HR). Depletion of mPges-1 in myeloid cells (LysM-$Cre^{+/-}$) did not significantly alter weight gain (D), SBP (E) and HR (F) in Pd-$1^{-/-}$/$Ldlr^{-/-}$ mice of both sexes fed a WD. Data are expressed as means ± SEMs (Mann-Whitney test, $*p < 0.05$, male, n= 5- 25 per group, female, n= 8- 20 per group).

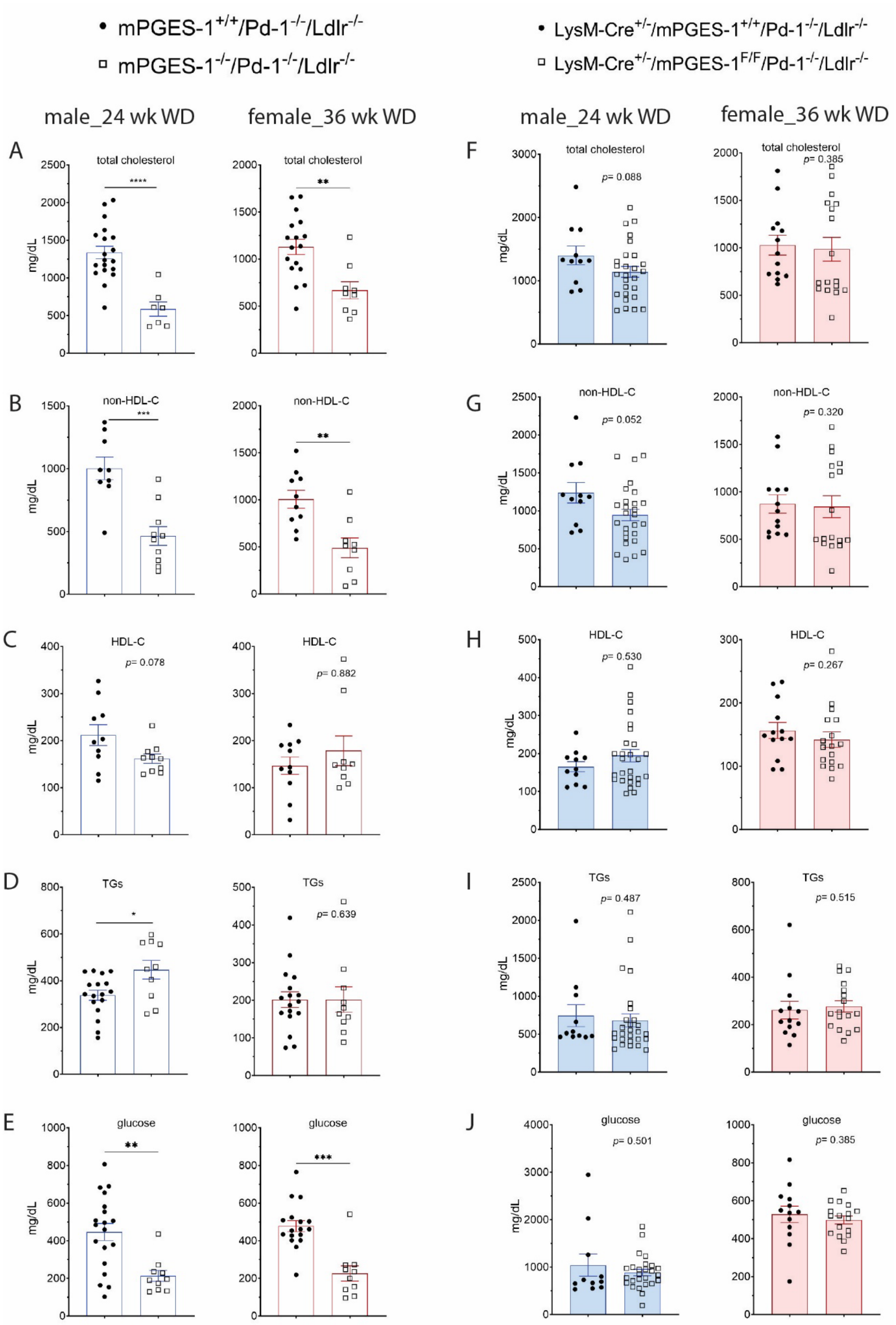

mPGES-1+/+/Pd-1-/-/Ldlr-/-
mPGES-1-/-/Pd-1-/-/Ldlr-/-
LysM-Cre+/-/mPGES-1+/+/Pd-1-/-/Ldlr-/-
LysM-Cre+/-/mPGES-1F/F/Pd-1-/-/Ldlr-/-
male_24 wk WD
female_36 wk WD
A
B
C
D
E
F
G
H
I
J
total cholesterol
non-HDL-C
HDL-C
TGs
glucose
mg/dL
p= 0.078
p= 0.882
p= 0.639
p= 0.088
p= 0.385
p= 0.052
p= 0.320
p= 0.530
p= 0.267
p= 0.487
p= 0.515
p= 0.501
p= 0.385

**Supplemental Figure S2. Depletion of mPges-1 globally significantly reduced plasma levels of total cholesterol, non-HDL cholesterol and glucose in Pd-1$^{-/-}$/Ldlr$^{-/-}$ mice fed a WD.** Plasma samples collected from male and female mice fed a WD for 24 weeks (male) and 36 weeks (female) were used to measure total cholesterol (A), (C) HDL cholesterol, (D) triglycerides and (E) glucose levels using biochemical kits following manufacturer's instructions. (B) Non-HDL cholesterol levels were calculated by subtracting HDL from total cholesterol. Depletion of mPges-1 in myeloid cells (LysM-Cre$^{+/-}$) did not significantly alter plasma levels of total cholesterol (F), non-HDL cholesterol (G), HDL cholesterol (H), TGs (I) and glucose (J) in Pd-1$^{-/-}$/Ldlr$^{-/-}$ mice of both sexes fed a WD. Data are expressed as means ± SEMs (Mann-Whitney test, **p*< 0.05, ***p*< 0.01, ****p*< 0.001 was considered significant, male, n= 9- 19 per group, female, n= 9- 18 per group)

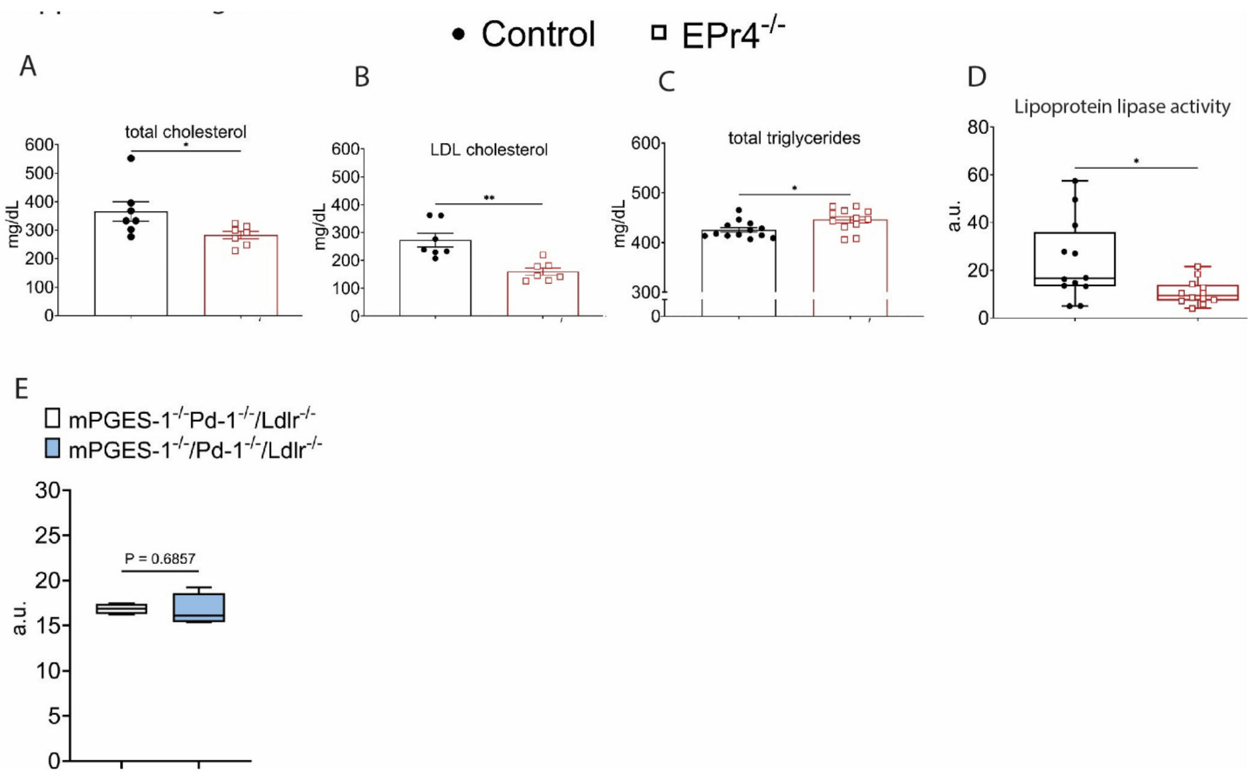

**Supplemental Figure S3. Depletion of Epr4 reduced plasma total and low-density lipoprotein cholesterol levels and lipoprotein lipase activity, while plasma triglycerides levels increased in female mice fed WD for 12 weeks.** Female mice were administered heparin (0.2U/ g body weight) by retroorbital route for 10 mins. Whole blood was centrifuged at 3000 g for 5 min to collect plasma to measure total and LDL cholesterol and triglyceride levels and lipoprotein lipase (LPL) activity following manufacture's instruction. (A-B) Plasma level of total cholesterol and LDL- cholesterol were decreased in EPr4$^{-/-}$ mice as compared to controls. Plasma level of triglycerides was elevated in the EPr4$^{-/-}$ mice (C). Depletion of Epr4$^{-/-}$ suppressed LPL activity in female mice after 12 weeks on a WD (D). $^{*}p< 0.05$, $^{**}p< 0.01$, n= 7- 13 per genotype. A Mann Whitney test (2-tailed) revealed a significant effect of Epr4 deletion on plasma total cholesterol, LDL cholesterol and triglyceride levels and LPL activity. (E) Deletion of mPges-1 globally did not significantly alter LPL in male Pd-1$^{-/-}$/Ldlr$^{-/-}$ mice fed a WD for 12 weeks. (Mann-Whitney test, $p> 0.05$ male, n= 4 per group). Data are expressed as means ± SEMs. Control- Ind-Cre$^{+/-}$/Pd-1$^{-/-}$/Ldlr$^{-/-}$ or EPr4$^{F/F}$/ Pd-1$^{-/-}$/Ldlr$^{-/-}$, EPr4$^{-/-}$- Ind-Cre$^{+/-}$/ EPr4$^{F/F}$/Pd-1$^{-/-}$/Ldlr$^{-/-}$.

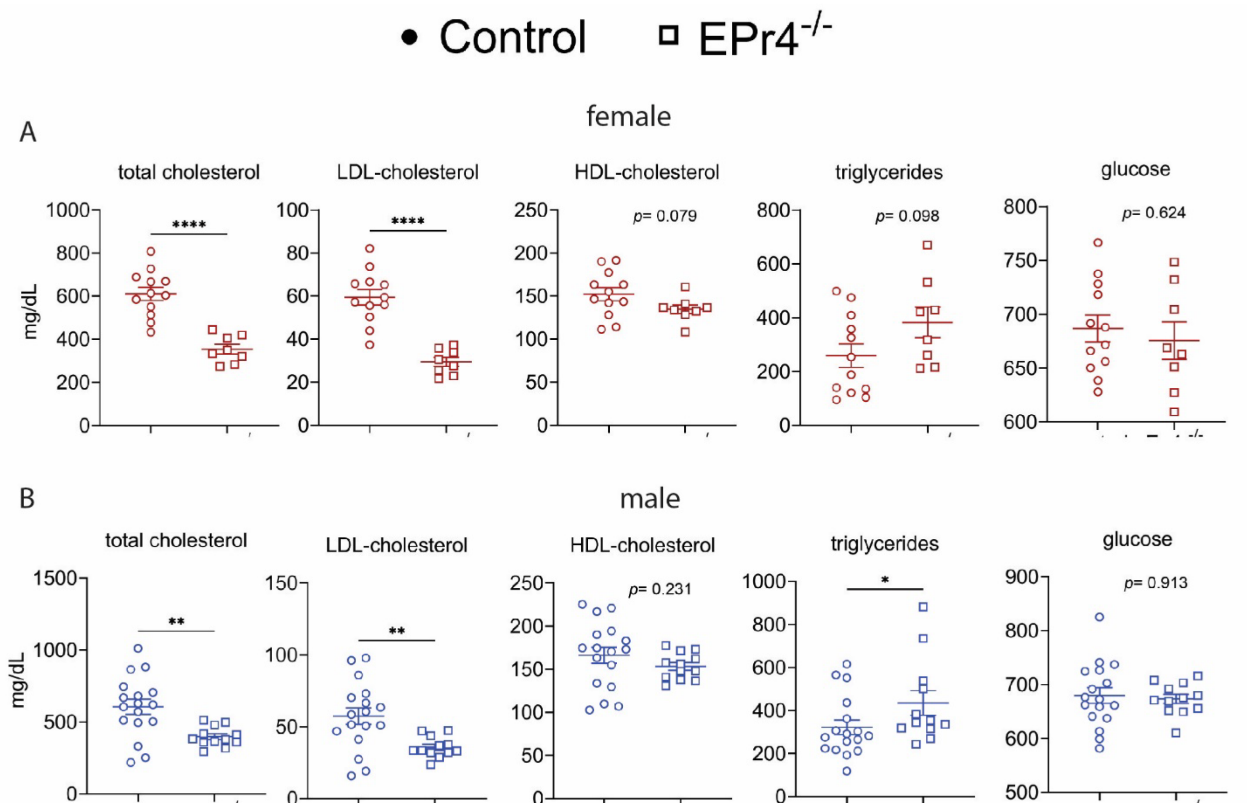


**Supplemental Figure S4. Depletion of EPr4 decreased total and LDL cholesterol, increased triglycerides in Pd-1$^{-/-}$/Ldlr$^{-/-}$ mice of both sexes fed a WD for 24 weeks.**

Plasma samples were obtained from whole blood after centrifugation at 3000g for 5 minutes at room temperature. Samples were kept at -80ºC until further analyses. Plasma total cholesterol, LDL-C, HDL-C, triglycerides, and glucose were determined using biochemical assay kits following manufacturers' instruction. Plasma levels Plasma levels of total cholesterol and LDL-cholesterol were decreased in EPr4$^{-/-}$ mice as compared to controls. By contrast, triglyceride levels were elevated in the EPr4$^{-/-}$ mice. $^{*}p< 0.05$, $^{**}p< 0.01$, female, n= 7- 12 per group, male, n= 12- 17 per group. A Mann Whitney test (2-tailed) revealed a significant effect of EPr4 deletion on total and LDL cholesterol and triglyceride levels. Data are expressed as means ± SEMs. Control- Ind-Cre$^{+/-}$/Pd-1$^{-/-}$/Ldlr$^{-/-}$ or EPr4$^{F/F}$/ Pd-1$^{-/-}$/Ldlr$^{-/-}$, EPr4$^{-/-}$- Ind-Cre$^{+/-}$/ EPr4$^{F/F}$/Pd-1$^{-/-}$/Ldlr$^{-/-}$.

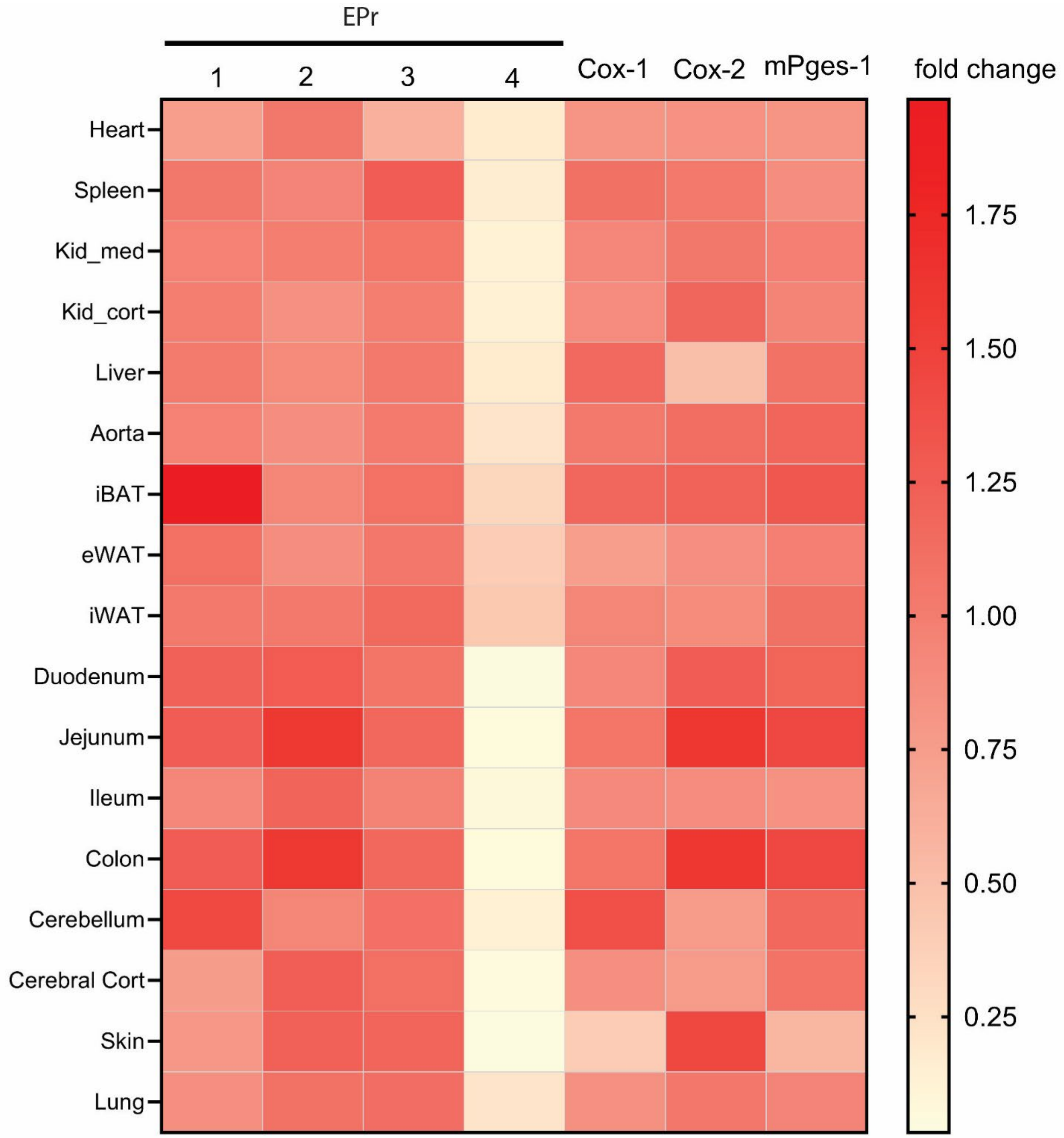


**Supplemental Figure S5. Tamoxifen-induced depletion of Epr4 in Pd-1$^{-/-}$/Ldlr$^{-/-}$ mice.** Heatmap of EPr4 gene expression in various tissues (EPr4$^{-/-}$ vs control). 17 different tissues from controls and EPr4 deficient mice were dissected for RNA extraction as detailed in the Supplemental Methods. EPr1, 2, 3, 4, Cox-1, Cox-2 and mPges-1 mRNA expression were measured by rt-qPCR. Results were expressed as fold change as compared to control mice. Kid med- kidney medulla, Kid cort- kidney cortex, iBAT- interscapular brown adipose tissue, eWAT- epididymal white adipose tissue, iWAT- inguinal white adipose tissue, cerebral cort- cerebral cortex. Control- Ind-Cre$^{+/-}$/Pd-1$^{-/-}$/Ldlr$^{-/-}$ or EPr4$^{F/F}$/ Pd-1$^{-/-}$/Ldlr$^{-/-}$, EPr4$^{-/-}$- Ind-Cre$^{+/-}$/ EPr4$^{F/F}$/Pd-1$^{-/-}$/Ldlr$^{-/-}$.

.

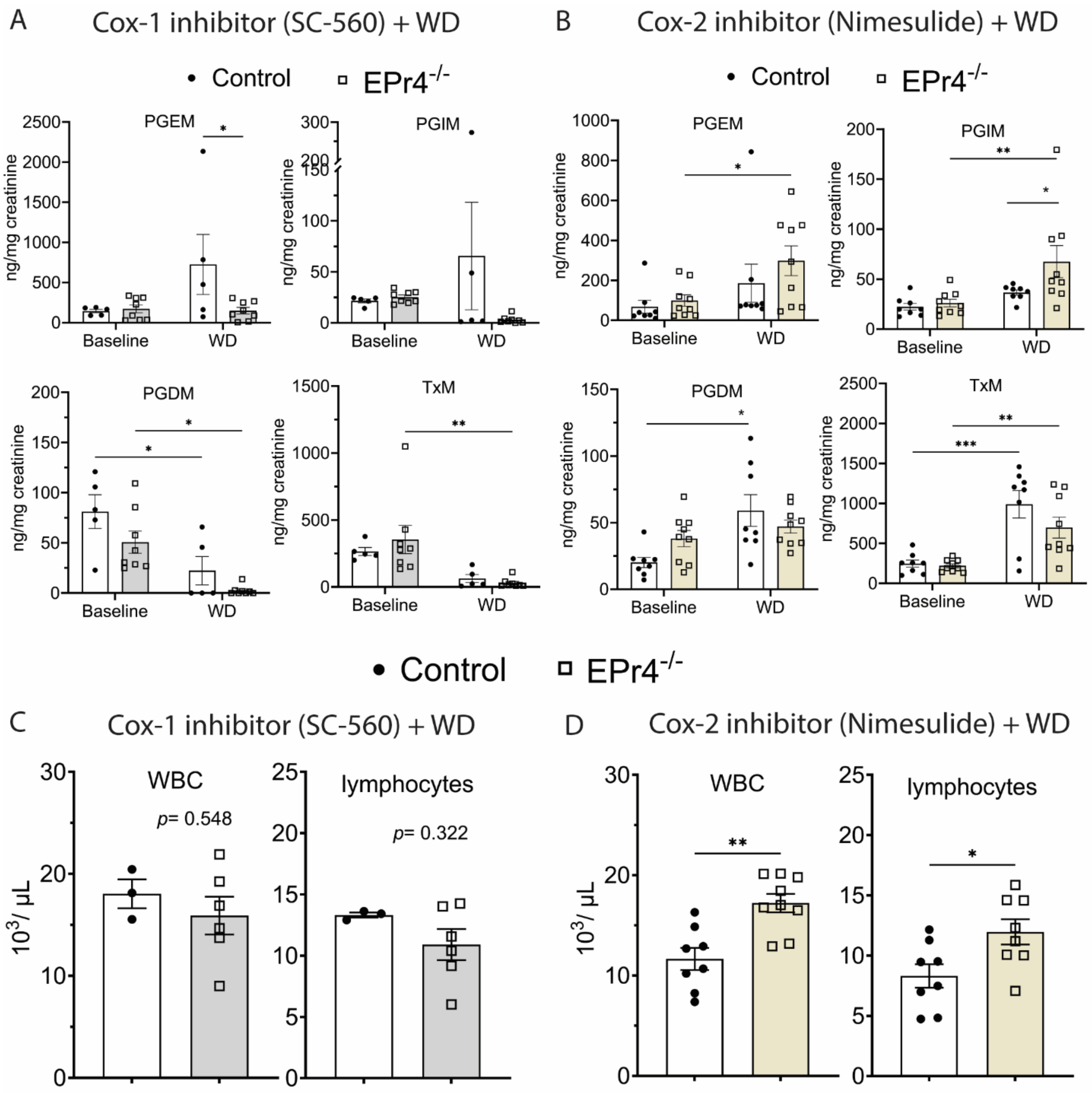


**Supplemental Figure S6. Urinary prostanoid metabolites of controls and EPr4$^{-/-}$ mice of both sexes fed a WD in conjunction with SC-560 (Cox-1 inhibitor) or Nimesulide (Cox-2 inhibitor).** Fasting (10am-5pm) urine samples from mice of both sexes were collected before and after feeding a WD for 6-8 weeks, and prostanoid metabolites (PGEM-7-hydroxy-5, 11-diketotetranorprostane-1, 16-dioic acid, PGIM- 2, 3-dinor 6-keto $PGF_{1\alpha}$, TxM- 2, 3-dinor $TxB_2$, tetranor PGDM- 11, 15-dioxo-9$_{\alpha}$-hydroxy-2, 3, 4, 5-tetranorprostan-1, 20-dioic acid, M-metabolite) were analyzed by LC/MS/MS as described in the Supplemental Methods. (A) SC-560 (15 mg/ kg BW, daily), a selective Cox-1 inhibitor was incorporated in WD and fed *ad libitum* to control and EPr4$^{-/-}$ mice. (B) Nimesulide (40 mg/ liter) was dissolved in drinking water and was available *ad libitum* to control and EPr4$^{-/-}$ mice. SC-560 significantly inhibited PGEM in EPr4$^{-/-}$ mice compared to control mice. (C-D) Whole blood collected from submandibular vein of mice on WD+ SC560 or WD+ nimesulide was used for complete blood count (CBC) analyses using the Sysmex Hematology Analyzer. CBC revealed no significant differences in WBC and lymphocyte numbers between control and EPr4$^{-/-}$ mice fed a WD + SC-560. On the contrary, WBC and lymphocytes numbers were significantly higher in EPr4$^{-/-}$ mice compared to control mice fed a WD+ nimesulide. For urinary prostanoid analyses, 2-way ANOVA showed that urinary prostanoid metabolites were significantly affected by SC-560 or nimesulide. Bonferroni multiple comparison test was used to test for significant differences between controls and EPr4$^{-/-}$ mice. Data are expressed as means ± SEMs. $^{*}p< 0.05$, $^{**}p< 0.01$, $^{***}p<0.001$. For SC-560 study, n= 3 females (2 controls, 1 EPr4$^{-/-}$), n= 10 males (3 controls, 7 EPr4$^{-/-}$). For nimesulide study, n= 10 females (6 controls, 4 EPr4$^{-/-}$), n= 7 males (2 controls, 5 EPr4$^{-/-}$). For CBC analyses, data are expressed as means ± SEMs, Mann-Whitney test, *$p< 0.05$, **$p< 0.01$ was considered significant. Control- Ind-Cre$^{+/-}$/Pd-1$^{-/-}$/Ldlr$^{-/-}$ or EPr4$^{F/F}$/ Pd-1$^{-/-}$/Ldlr$^{-/-}$, EPr4$^{-/-}$- Ind-Cre$^{+/-}$/ EPr4$^{F/F}$/Pd-1$^{-/-}$/Ldlr$^{-/-}$. M- metabolite.

A

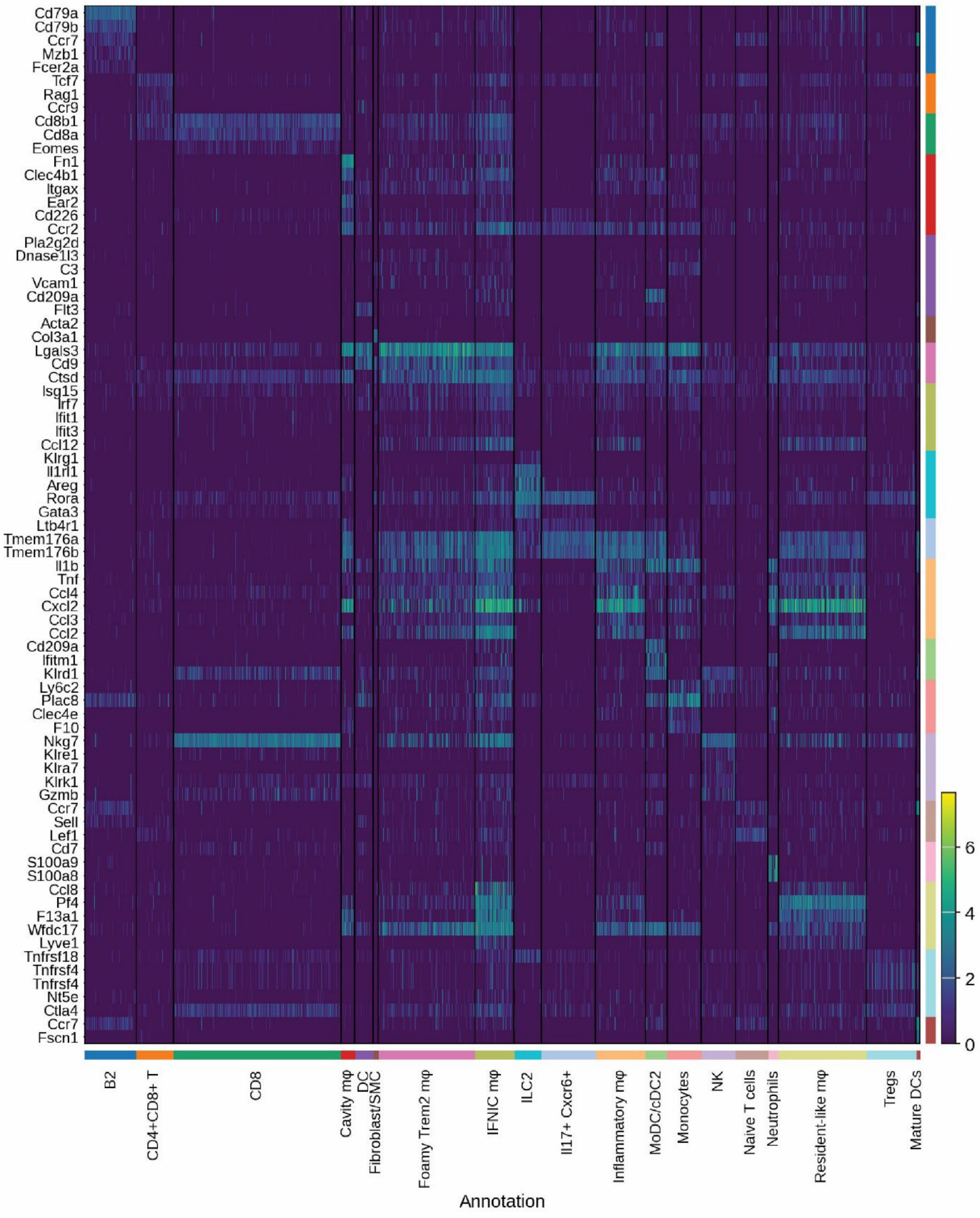

Cd79a
Cd79b
Ccr7
Mzb1
Fcer2a
Tcf7
Rag1
Ccr9
Cd8b1
Cd8a
Eomes
Fn1
Clec4b1
Itgax
Ear2
Cd226
Ccr2
Pla2g2d
Dnase1l3
C3
Vcam1
Cd209a
Flt3
Acta2
Col3a1
Lgals3
Cd9
Ctsd
Isg15
Irf7
Ifit1
Ifit3
Ccl12
Klrg1
Il1rl1
Areg
Rora
Gata3
Ltb4r1
Tmem176a
Tmem176b
Il1b
Tnf
Ccl4
Cxcl2
Ccl3
Ccl2
Cd209a
Ifitm1
Klrd1
Ly6c2
Plac8
Clec4e
F10
Nkg7
Klre1
Klra7
Klrk1
Gzmb
Ccr7
Sell
Lef1
Cd7
S100a9
S100a8
Ccl8
Pf4
F13a1
Wfdc17
Lyve1
Tnfrsf18
Tnfrsf4
Tnfrsf4
Nt5e
Ctla4
Ccr7
Fscn1
6
4
2
0
B2
CD4+CD8+ T
CD8
Cavity mφ
DC
Fibroblast/SMC
Foamy Trem2 mφ
IFNIC mφ
ILC2
Il17+ Cxcr6+
Inflammatory mφ
MoDC/cDC2
Monocytes
NK
Naive T cells
Neutrophils
Resident-like mφ
Tregs
Mature DCs
Annotation

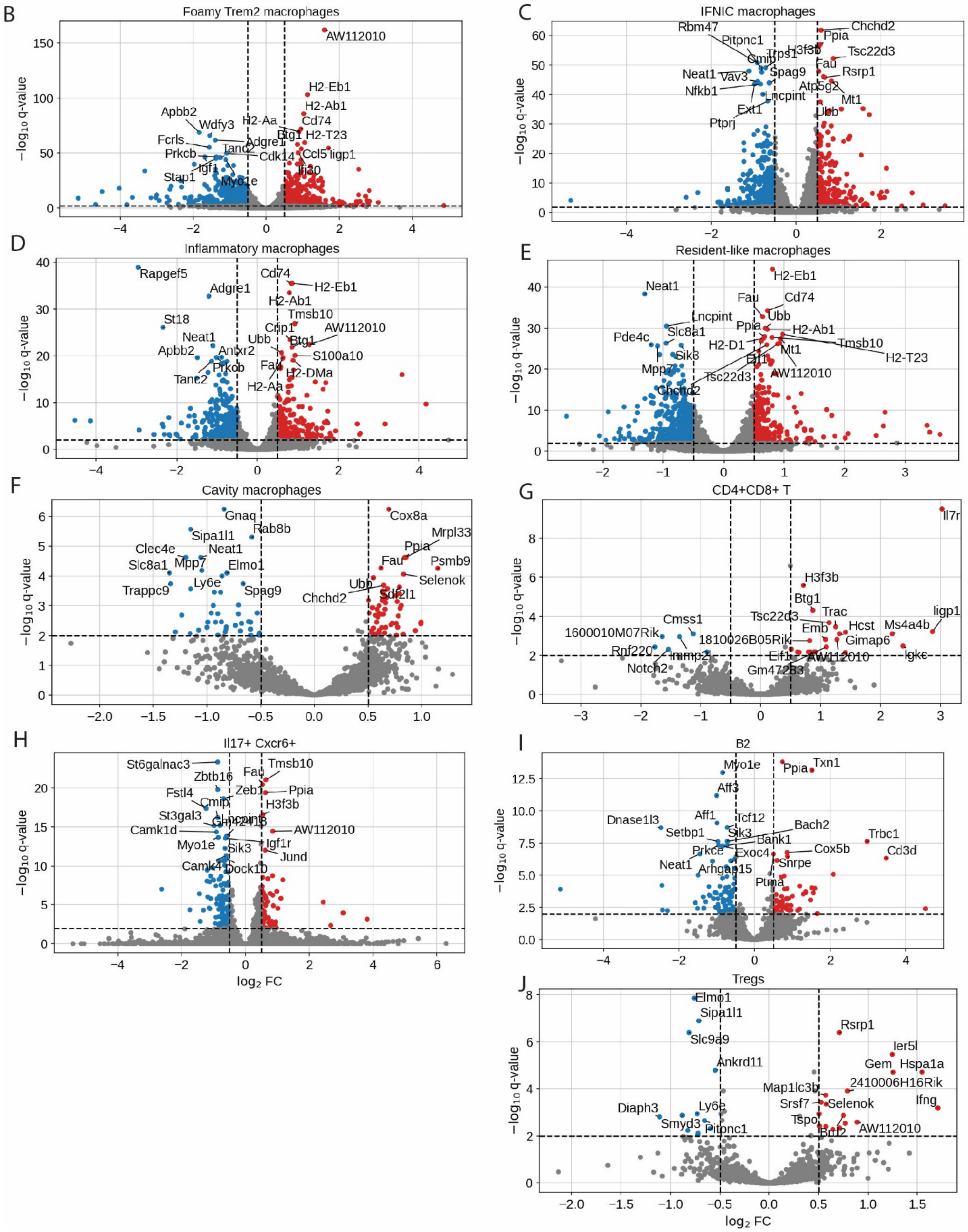

B
Foamy Trem2 macrophages
AW112010
H2-Eb1
H2-Ab1
Cd74
H2-Aa
Btg1
H2-T23
Apbb2
Wdfy3
Adgre1
Fcrls
Tanc2
Prkcb
Cdk14
Ccl5
Iigp1
Igf1
Stap1
Myo1e
Ifi30
C
IFNIC macrophages
Rbm47
Pitpnc1
Chchd2
Ppia
Tsc22d3
Fau
Rsrp1
Atp5g2
Mt1
Ubb
Neat1
Vav3
Spag9
Nfkb1
Lncpint
Ext1
Ptprj
D
Inflammatory macrophages
Rapgef5
Adgre1
St18
Neat1
Apbb2
Anxr2
Prkcb
Tanc2
Cd74
H2-Eb1
H2-Ab1
Tmsb10
AW112010
Btg1
Ubb
S100a10
Fau
H2-DMa
H2-Aa
E
Resident-like macrophages
Neat1
H2-Eb1
Fau
Cd74
Lncpint
Ubb
Slc8a1
Ppia
H2-Ab1
Pde4c
H2-D1
Tmsb10
Sik3
Mt1
H2-T23
Mpp7
Tsc22d3
AW112010
Chchd2
F
Cavity macrophages
Gnaq
Cox8a
Rab8b
Sipa1l1
Mrpl33
Clec4e
Neat1
Ppia
Slc8a1
Mpp7
Fau
Psmb9
Elmo1
Selenok
Ly6e
Trappc9
Spag9
Chchd2
Ubb
Sdf2l1
G
CD4+CD8+ T
Il7r
H3f3b
Btg1
Trac
Cmss1
Tsc22d3
Hcst
Ms4a4b
Iigp1
1600010M07Rik
1810026B05Rik
Emb
Gimap6
Rnf220
Eif1
AW112010
Igkc
Notch2
Gm47283
H
Il17+ Cxcr6+
St6galnac3
Tmsb10
Fau
Zbtb16
Zeb1
Ppia
Fstl4
Cmip
H3f3b
St3gal3
Camk1d
AW112010
Myo1e
Igf1r
Sik3
Jund
Camk4
Dock10
I
B2
Myo1e
Ppia
Txn1
Aff3
Aff1
Tcf12
Dnase1l3
Bach2
Setbp1
Sik3
Trbc1
Bank1
Prkce
Exoc4
Cox5b
Cd3d
Neat1
Snrpe
Arhgap15
Ptma
J
Tregs
Elmo1
Sipa1l1
Rsrp1
Slc9a9
Ier5l
Gem
Hspa1a
Ankrd11
2410006H16Rik
Map1lc3b
Srsf7
Selenok
Ifng
Diaph3
Ly6e
Smyd3
Tspo
Pitpnc1
Btf2
AW112010
$-\log_{10}$ q-value
$\log_2$ FC

**Supplemental Figure S7. scRNA analysis of $CD45^+$ cells isolated from atherosclerotic lesions of female mice fed a WD for 12 weeks**. (A) Heatmap of the marker genes in each cluster selected as unique genes used for identification of each cluster. (B-J) Volcano plots of differential expressed genes (DEGs) of macrophages and T cells in atherosclerotic lesions in female mice fed a WD for 12 weeks. (B) foamy Trem2 Mϴ, (C) IFNIC Mϴ, (D) inflammatory Mϴ, (E) resident-like Mϴ, (F) cavity Mϴ, (G) $CD4^+CD8^+$ T cells, (H) $Il17^+Cxcr6^+$ cells, (I) B2 cells, (J) Tregs cells. More upregulated (red) and downregulated (blue) genes in different macrophage cell types (B, C, D, E) were identified in $EPr4^{-/-}$ cells as compared to T cells (G, H, J) and B cells (I).

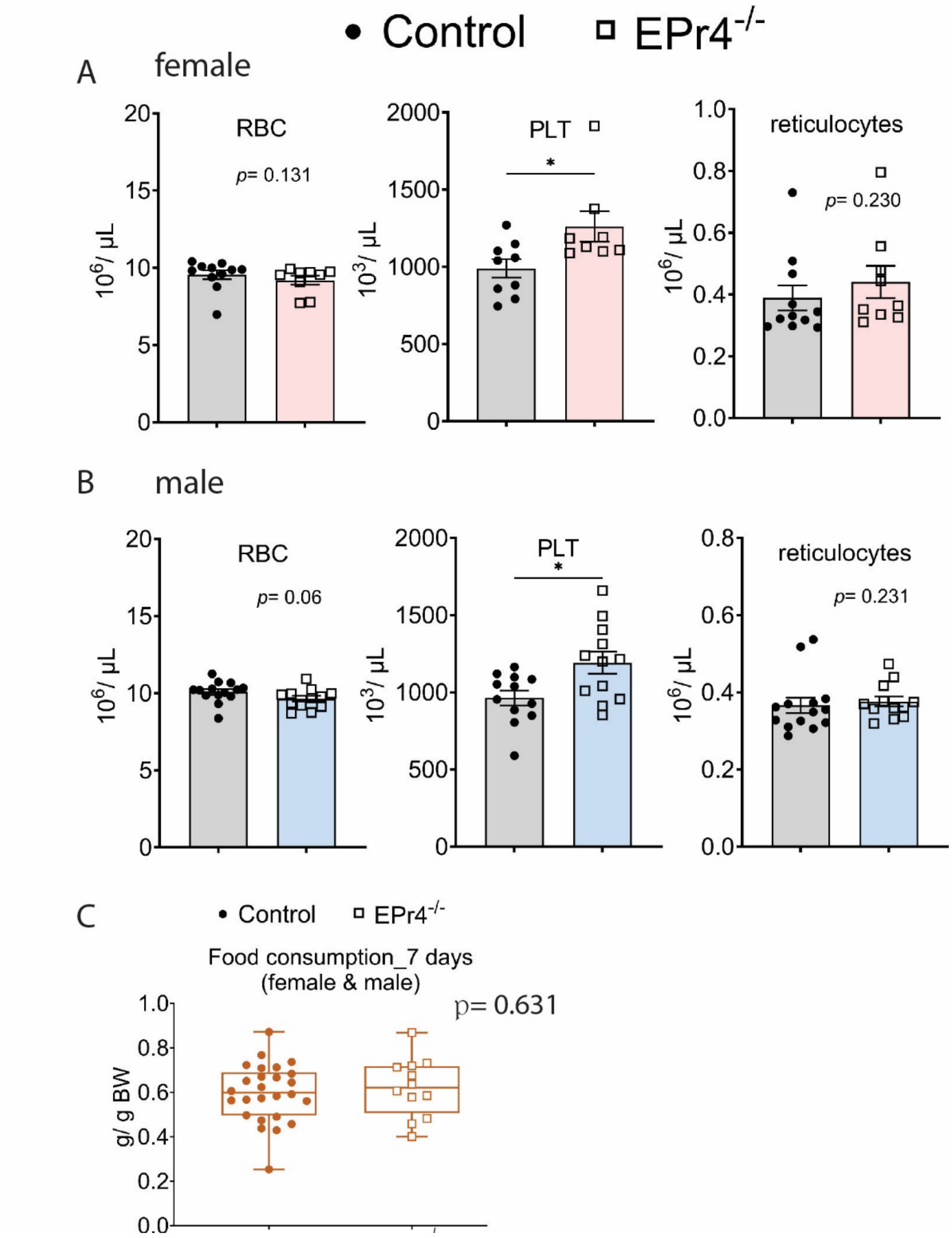


**Supplemental Figure S8. Platelet count (PLT) was significantly increased in EPr4$^{-/-}$ mice, depletion of Epr4 did not significantly alter WD consumption in Pd-1$^{-/-}$/Ldlr$^{-/-}$ mice.** (A & B) Complete blood count (CBC) was performed using a Sysmex Hematology Analyzer. CDC analyses revealed a significant difference in the number of platelets (PLT) between controls and EPr4 deficient Pd-1$^{-/-}$/Ldlr$^{-/-}$ mice of both sexes fed a WD for 12 weeks. A Mann Whitney test (2-

tailed) revealed a significant effect of EPr4 genotype on PLT. *$p< 0.05$, n= 7- 12 per group. (C) The amount of WD consumed by mice housed individually was estimated by weighing the food for 7 days. A Mann Whitney test (2-tailed) revealed no significant difference in WD consumption between controls and $EPr4^{-/-}$ mice. Data are expressed as means ± SEMs. $p>0.05$, n= 12- 26 per group. BW- body weight. Control- Ind-$Cre^{+/-}$/Pd-$1^{-/-}$/$Ldlr^{-/-}$ or $EPr4^{F/F}$/ Pd-$1^{-/-}$/$Ldlr^{-/-}$, $EPr4^{-/-}$- Ind-$Cre^{+/-}$/ $EPr4^{F/F}$/Pd-$1^{-/-}$/$Ldlr^{-/-}$.

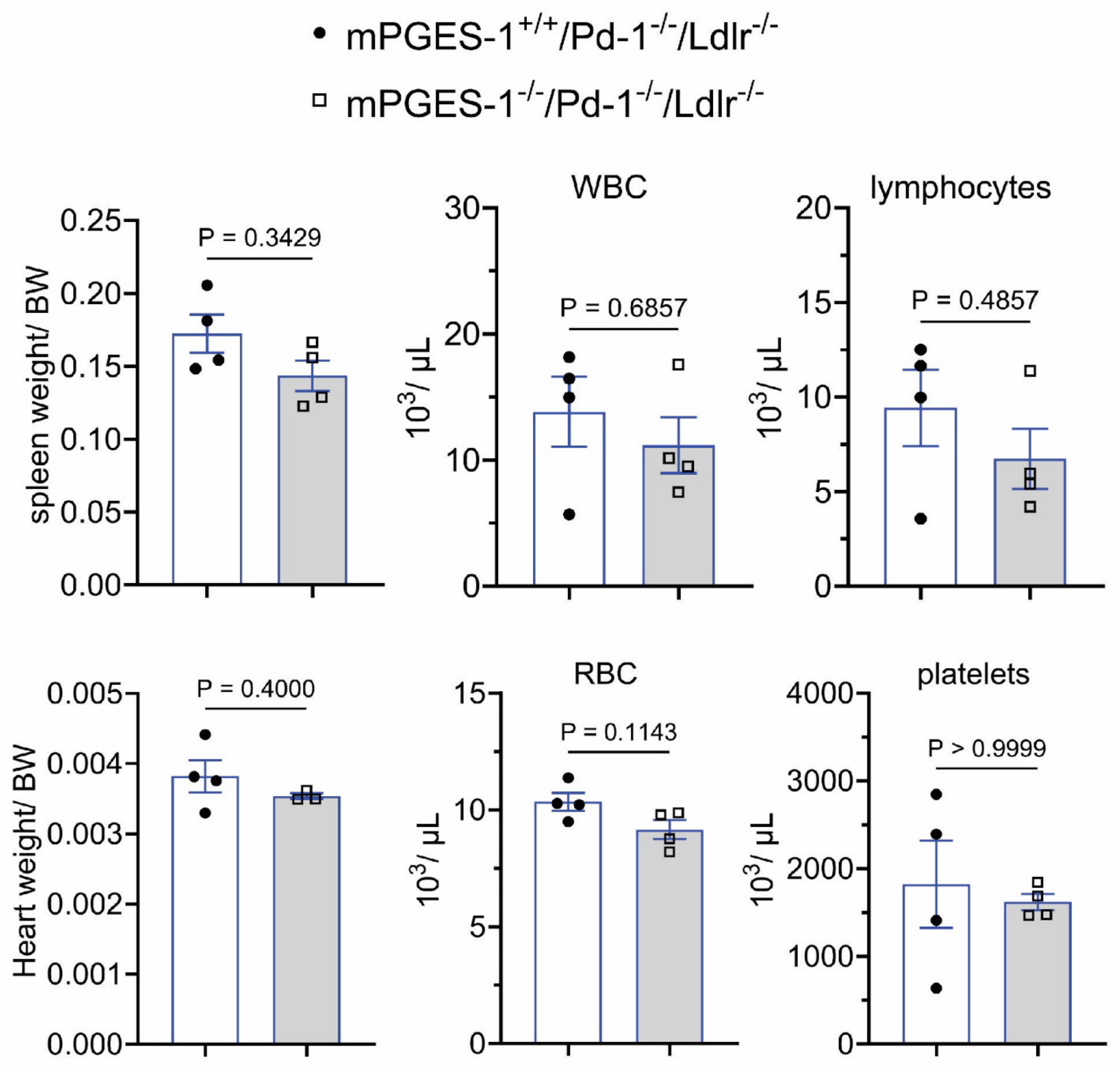

**Supplemental Figure S9. Depletion of mPges-1 did not significantly affect heart and spleen weights, numbers of WBC, RBC, lymphocytes, and platelets in Pd-1$^{-/-}$/Ldlr$^{-/-}$ mice.**

Male mice on a WD for 12 weeks were studied in this experiment. (A & B) Heart and spleen were weighed and normalized with body weight. Whole blood was collected in heparin coated tube and complete blood count (CBC) was performed using a Sysmex Hematology Analyzer. A Mann Whitney test (2-tailed) revealed no significant effects of mPges-1 deletion on heart and spleen weights, WBC, RBC, lymphocyte and platelet numbers in whole blood. Data are expressed as means ± SEMs. $P > 0.05$, n= 3-4 per group. BW- body weight.

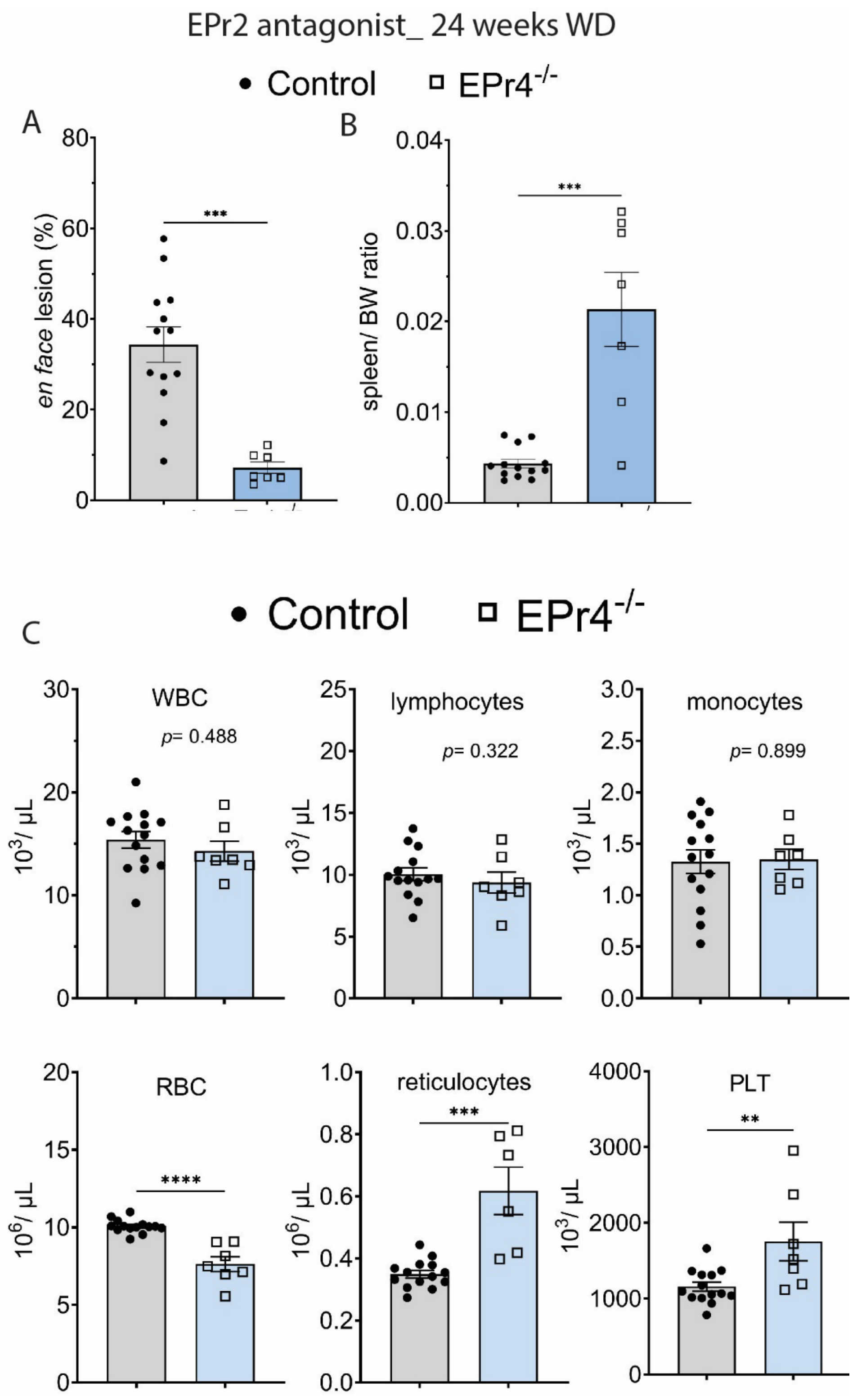
EPr2 antagonist_ 24 weeks WD
Control
EPr4$^{-/-}$
A
en face lesion (%)
80
60
40
20
0
***
B
spleen/ BW ratio
0.04
0.03
0.02
0.01
0.00
***
C
Control
EPr4$^{-/-}$
WBC
p= 0.488
$10^3$/ µL
30
20
10
0
lymphocytes
p= 0.322
$10^3$/ µL
25
20
15
10
5
0
monocytes
p= 0.899
$10^3$/ µL
3.0
2.5
2.0
1.5
1.0
0.5
0.0
RBC
****
$10^6$/ µL
20
15
10
5
0
reticulocytes
***
$10^6$/ µL
1.0
0.8
0.6
0.4
0.2
0.0
PLT
**
$10^3$/ µL
4000
3000
2000
1000
0

**Supplemental Figure S10. Epr2 blockade did not confer additional restrain on *en face* aortic lesions in male EPr4 deficient Pd-1$^{-/-}$/Ldlr$^{-/-}$ male mice fed a WD for 24 weeks.** Aortic atherosclerotic lesion burden, represented by the percentage of lesion area to total aortic area, was quantified by *en face* analysis of aortas stained with Sudan IV from mice fed a WD. EPr2 antagonist was incorporated into WD and fed to control and EPr4$^{-/-}$ Pd-1$^{-/-}$/ Ldlr$^{-/-}$ mice for 24 weeks. (A) EPr2 blockade did not confer additional restrain on the accumulation of aortic plaques in EPr4 deficient Pd-1$^{-/-}$/ Ldlr$^{-/-}$ mice fed a WD for 24 weeks. (B) Spleen size normalized to body weight was significantly higher in EPr4$^{-/-}$ mice as compared to controls mice fed WD+ EPr2 antagonist. (C) Complete blood count (CBC) was performed using a Sysmex Hematology Analyzer. CDC analyses revealed no significant differences in the numbers of WBC, lymphocytes and monocytes between controls and EPr4 deficient Pd-1$^{-/-}$/Ldlr$^{-/-}$ fed a WD+ EPr2 antagonist. RBC decreased while reticulocytes and PLT increased in Epr4 deficient mice. A Mann Whitney test (2-tailed) revealed a significant effect of EPr4 genotype on aortic *en face* lesion burden and spleen size. Data are expressed as means ± SEMs. ***$p$< 0.001, n= 6- 14 per group. Control- Ind-Cre$^{+/-}$/Pd-1$^{-/-}$/Ldlr$^{-/-}$ or EPr4$^{F/F}$/ Pd-1$^{-/-}$/Ldlr$^{-/-}$, EPr4$^{-/-}$- Ind-Cre$^{+/-}$/ EPr4$^{F/F}$/Pd-1$^{-/-}$/Ldlr$^{-/-}$.

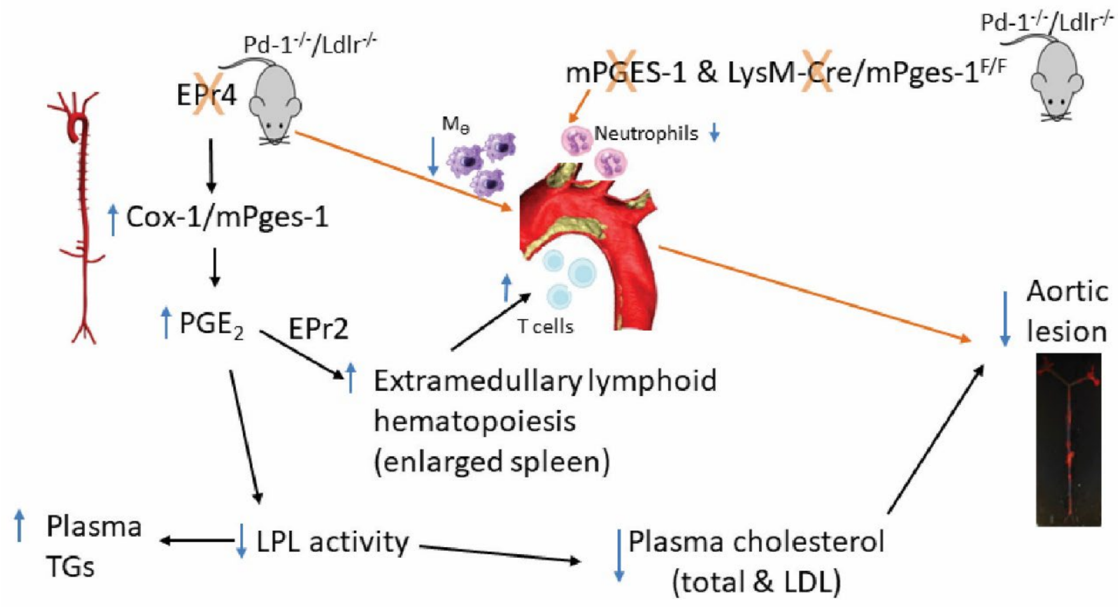


**Supplemental Figure S11. Schematic illustrating the mechanism of mPGES-1/ $PGE_2$/ EPr4 signaling pathways in promoting macrophages infiltration and atherosclerosis.** Depletion of mPges-1 globally, specifically in myeloid cells, or Epr4 globally restrains atherogenesis in Pd-1$^{-/-}$ / Ldlr$^{-/-}$ mice. These effects are attributable to the roles of $PGE_2$/ EPr4 signaling in promoting chemotaxis of myeloid cells to inflamed sites during disease progression. Depletion of mPges-1 globally reduced aortic neutrophils in male mice. Upregulation of $PGE_2$ biosynthesis in EPr4 deficient mice suppresses lipoprotein lipase activity (LPL), coincident with the reduction in plasma cholesterol (total and LDL) and increase in triglyceride levels. Additionally, $PGE_2$ via EPr2 promotes extramedullary lymphoid hematopoiesis in the spleen. Consequently, T cells expanded in atherosclerotic plaques.
