## Supplemental methods for "Disruption of the PGE_2_ synthesis / response pathway restrains atherogenesis in programmed cell death-1 (Pd-1) deficient hyperlipidemic mice"

**Short title for running head: mPGES-1/ $PGE_2$/ Ep4r signaling in Pd-1$^{-/-}$/ Ldlr$^{-/-}$ mice**

[1, 2#]Emanuela Ricciotti, PhD; [1#]Soon Yew Tang, PhD; [1]Antonijo Mrčela, PhD; [1]Ujjalkumar S. Das, PhD; [1]Ronan Lordan, PhD; [1]Robin Joshi, PhD; [1]Soumita Ghosh, PhD; [1]Justin Aoyama, BSc; [1]Ryan McConnell, BSc; [1]Jianing Yang, PhD; [1,3]Gregory R. Grant, PhD; [1,4]*Garret A. FitzGerald, MD

[1]From the Institute for Translational Medicine and Therapeutics, Perelman School of Medicine, [2]the Department of Systems Pharmacology and Translational Therapeutics, the [3]Department of Genetics, and the [4]Department of Medicine Perelman School of Medicine, University of Pennsylvania.

*Address for correspondence: Garret A. FitzGerald, Institute for Translational Medicine and Therapeutics, Perelman School of Medicine, 10-110 Smilow Center for Translational Research, 3400 Civic Center Blvd, Bldg 421, University of Pennsylvania, Philadelphia, PA 19104-5158. Fax: 215-573-9135 Tel: 215-898-1184

### These authors contributed equally.

#### Supplemental methods

##### Genotyping protocol for Ind-Cre$^{+/-}$, LysM-Cre$^{+/-}$, mPges-1, mPges-1 Flox/Flox, EPr4 Flox/Flox, Pd-1, and Ldlr.

Genotypes were determined by polymerase chain reaction (PCR) with primers specific for each allele:

Ind-Cre$^{+/-}$: forward primer (FP)- 5'-GCATTGCTGTCACTTGGTCG-3'; reverse primer (RP)- 5'-GCATTGCTGTCACTTGGTCGT-3'; Cre mutant allele: 250 bp. PCR program: 1) 94ºC, 1min; 2) 94ºC, 30sec; 3) 50ºC, 30sec; 4) 72ºC, 60sec; 5) repeat steps 2-4 for 30 cycles; 6) 72ºC, 7min; 7) 10ºC, hold.

LysM-Cre$^{+/-}$: FP- 5'-CCCAGAAATGCCAGATTACG-3'; RP1- 5'-CTTGGGCTGCCAGAATTTCTC-3'; FP2- 5'-TTACAGTCGGCCAGGCTGAC-3'; wild-type allele: 350 bp; Cre mutant allele: 700 bp. PCR program: 1) 94ºC, 1min; 2) 94ºC, 30sec; 3) 58ºC, 30sec; 4) 72ºC, 60sec; 5) repeat steps 2-4 for 35 cycles; 6) 72ºC, 7min; 7) 10ºC, hold.

mPges-1: S1- 5'-TCCCAGGTGTTGGGATTTAGACG-3'; S2- 5'-TAGGTGGCTGTACTGTTTGTTGC -3'; S3- 5'-ACTCCAGTACTGAGCCAGCTGC -3'; PGK- 5'-TGCTACTTCCATTTGTCACGTCC -3'; wild-type allele: 400 bp, mutant allele: 800 bp. PCR program: 1) 94ºC, 3min; 2) 94ºC, 50sec; 3) 64ºC, 55sec; 4) 72ºC, 1min 5) repeat steps 2-4 for 5 cycles; 6) 94ºC, 50sec; 7) 62ºC, 55sec; 8) 72ºC, 1min 10sec 9) repeat steps 6-8 for 10 cycles; 10) 94ºC, 50sec; 11) 59ºC, 55sec; 12) 72ºC, 1min 10sec 13) repeat steps 10-12 for 10 cycles; 14) 94ºC, 50sec; 15) 56ºC, 55sec; 16) 72ºC, 1min 10sec 17) repeat steps 14-16 for 10 cycles; 18) 72ºC, 10min; 19) 10ºC, hold.

mPges-1 Flox/Flox: FP- 5'-AAGGTTGATGGTGCACGTCTA -3'; RP- 5'-TTTGTTTGGCCTCACTCTACA -3'; wild-type allele: 279 bp, floxed allele: 399 bp. PCR program: 1) 93ºC, 2min; 2) 93ºC, 10sec; 3) 55ºC, 40sec; 4) 65ºC, 1min 5) repeat steps 2-4 for 40 cycles; 6) 65ºC, 5min; 7) 10ºC, hold.

EPr4 Flox/Flox: FP1- 5'-GGCGGGATCAGTTAGATGG -3'; RP- 5'-GTGAAGCGAGTCCTTAGGC -3'; wild-type: 250 bp; floxed allele: 340 bp. PCR program: 1) 94ºC, 2min; 2) 94ºC, 20sec; 3) 65ºC, 15sec; 4) 68ºC, 10sec 5) repeat steps 2-4 for 10 cycles; 6) 94ºC, 15sec; 7) 60ºC, 15sec; 8) 72ºC, 10sec; 9) repeat steps 6-8 for 28 cycles; 10) 72ºC, 2min; 11) 10ºC, hold.

Pd-1: FP-5'-TCCTGCCAAACCTTGTAGTCA-3'; RP1- 5'-ACAACACAGGGTAGGCATGTAGCA-3'; neo primer- 5'-GCTAGCCAACCAGAAGTCTAA -3'; wild-type allele: 234 bp; mutant allele: 325 bp. PCR program: 1) 94ºC, 2min; 2) 94ºC, 60sec; 3) 60ºC, 60sec; 4) 72ºC, 1min 30sec 5) repeat steps 2-4 for 35 cycles; 6) 72ºC, 5min; 7) 10ºC, hold.

Ldlr: FP-5'-AAGACGTGCTCCCAGGATG-3'; RP- 5'-CGTGCTCCTCATCTGACTTGT-3'; neo primer- 5'-ATCGCCTTCTTGACGAGTTC-3'; wild-type allele: 383 bp; mutant allele: 550 bp. PCR program: 1) 90ºC, 1min; 2) 94ºC, 30sec; 3) 61ºC, 30sec; 4) 72ºC, 1min 5) repeat steps 2-4 for 34 cycles; 6) 72ºC, 7min; 7) 10ºC, hold.

EmeraldAmp GT PCR Master Mix (2x Premix) was used for all genotyping protocols.

**Generation of mPges-1-deficient, myeloid-specific mPges-1 deficient, Ind-Cre$^{+/-}$/ EPr4$^{F/F}$ knockout mice on Pd-1$^{-/-}$/ Ldlr$^{-/-}$ background**

All the mice used in the current study were on a low-density lipoprotein receptor deficient background (Ldlr$^{-/-}$ mice on a C57BL/6J background were purchased from the Jackson Laboratory) unless otherwise stated. Male Pd-1$^{-/-}$ mice were mated with female Ldlr$^{-/-}$ mice to obtain Pd-1$^{-/-}$/ Ldlr$^{-/-}$ mice after a second round of Pd-1$^{+/-}$/ Ldlr$^{+/-}$ intercrossing. Similarly, male mPges-1$^{-/-}$ mice were mated with female Pd-1$^{-/-}$/ Ldlr$^{-/-}$ mice to obtain mPges-1$^{+/-}$/ Pd-1$^{-/-}$/ Ldlr$^{-/-}$

mice. Intercrossing of mPges-$1^{+/-}$/ Pd-$1^{-/-}$/ Ldlr $^{-/-}$ allowed us to select mPges-$1^{+/+}$ Pd-$1^{-/-}$/ Ldlr $^{-/-}$ as controls and mPges-$1^{-/-}$/ Pd-$1^{-/-}$/ Ldlr $^{-/-}$ as global mPges-$1^{-/-}$ mice. Same breeding strategies were followed to obtain myeloid-specific mPges-1-deficient mice using LysM-Cre$^{+/-}$ and mPges-$1^{F/F}$ mice (1) on Pd-$1^{-/-}$/ Ldlr$^{-/-}$ background. Male LysM-Cre$^{+/-}$/ mPges-$1^{F/+}$/ Pd-$1^{-/-}$/ Ldlr$^{-/-}$ mice were mated with female mPges-$1^{F/+}$/ Pd-$1^{-/-}$/ Ldlr$^{-/-}$ to yield LysM-Cre$^{+/-}$/ mPges-$1^{+/+}$/ Pd-$1^{-/-}$/ Ldlr$^{-/-}$ as controls and LysM-Cre$^{+/-}$/ mPges-$1^{F/F}$/ Pd-$1^{-/-}$/ Ldlr$^{-/-}$ as myeloid-specific mPges-$1^{-/-}$ mice. For global EPr4 deficient mice on Pd-$1^{-/-}$/ Ldlr$^{-/-}$ backgrounds, male Ind-Cre$^{+/-}$ mice were mated with female EPr4$^{F/F}$ mice to generate Ind-Cre$^{+/-}$/ EPr4$^{F/+}$ mice. The male Ind-Cre$^{+/-}$/ EPr4$^{F/+}$mice were then mated with female Pd-$1^{-/-}$/ Ldlr$^{-/-}$ mice to select for Ind-Cre$^{+/-}$/ EPr4$^{F/+}$/ Pd-$1^{+/-}$/ Ldlr$^{+/-}$ mice, and at the same time female EPr4$^{F/F}$ mice were mated with male d-$1^{-/-}$/ Ldlr$^{-/-}$ mice to generate EPr4$^{F/+}$/ Pd-$1^{+/-}$/ Ldlr$^{+/-}$ mice. Inter-crossing of EPr4$^{F/+}$/ Pd-$1^{+/-}$/ Ldlr$^{+/-}$ mice allowed us to select EPr4$^{F/+}$/ Pd-$1^{-/-}$/ Ldlr$^{-/-}$ mice for further breeding with Cre mice. Male Ind-Cre$^{+/-}$/ EPr4$^{F/+}$/ Pd-$1^{-/-}$/ Ldlr$^{-/-}$ mouse lines were crossed with female EPr4$^{F/+}$/ Pd-$1^{-/-}$/ Ldlr$^{-/-}$ to generate Ind-Cre$^{+/-}$/ Pd-$1^{-/-}$/ Ldlr$^{-/-}$ or EPr4$^{F/F}$/ Pd-$1^{-/-}$/Ldlr$^{-/-}$ (as control mice) and Ind-Cre$^{+/-}$/ EPr4$^{F/F}$/ Pd-$1^{-/-}$/ Ldlr$^{-/-}$ (as global EPr4$^{-/-}$ mice).

**Atherogenesis study**

Mice on Ldlr$^{-/-}$ background of both sexes were fed a Western Diet (WD, 21% fat, 1.5% cholesterol, 12079B, Research Diets, New Brunswick, NJ) from eight or ten weeks of age for 12, 24, or 36 weeks. Body weights were recorded, and urine samples were collected from the mice before and after the WD feeding.

**Blood pressure and heart rate measurement using tail-cuff system for Western diet (WD) fed mice**

Systolic blood pressure (SBP) and heart rate (HR) were measured in conscious mice on Ldlr$^{-/-}$ background using a computerized non-invasive tail-cuff system (Visitech Systems, Apex, NC) as described.(2) SBP and HR were recorded once each day from 8am to 11am for three consecutive days after two days of acclimatization on the system. Average SBP and HR were reported.

**Preparation of mouse aortas and *en face* quantification of atherosclerosis**

At the end of 12, 24 or 36 weeks on a WD, mice on Ldlr$^{-/-}$ background were euthanized between 8am- 11am by $CO_2$ overexposure in no particular order with respect to sex or phenotype. Mouse aorta was perfused *in situ* with ice-cold phosphate-buffered solution (PBS) and dissected from the aortic arch to the iliac bifurcation. The dissected aortas were fixed in Prefer fixative at 4ºC for at least 12 h. The extent of atherosclerosis (Phase 3 Imaging Systems, Glen Mills, PA) was determined by the *en face* methods and by assessment of aortic root lesion burden. For analysis of aortic plaques, the area of plaques stained positive with Sudan IV was expressed as percentage of the total area of the entire aorta.

For EPr2 antagonist study, PF-04418948 (10 mg/Kg BW, #HY-18966, MCE) was incorporated in WD. Controls and EPr4$^{-/-}$ male mice were fed this diet for 24 weeks and sacrificed for aortic lesion quantification by *en face* method as described previously.

**Assessment of aortic root lesion burden by haemotoxylin and eosin staining**

Optimal cutting temperature compound (OCT)-fixed aortic roots (n=5- 6) from each group were serially sectioned into 5 µm thickness and mounted on Superfrost Plus slides for analysis of tissue morphology by H&E staining. Eight sections (each 50 µm apart) were collected starting from the appearance of the tricuspid valve. Due to technical challenges in getting quality tissues sections, not all sections intended for comparison were suitable for analysis. Instead, an investigator blinded to genotype randomly selected two sufficient quality sections on each slide of the samples, which were used to assess aortic root lesion burden using Image J software. Area of aortic root lesions was expressed as percentage of the total area of the vessel minus the luminal area.

**Immunohistochemical examination of lesion morphology**

Superfrost Plus slides with aortic root sections were fixed in ice-cold acetone for 15 min at -20ºC. Before incubating with primary antibodies, sections were consecutively incubated with 3% $H_2O_2$ to remove endogenous peroxidase, 10% normal serum blocking solution (dependent on host secondary antibody) diluted with Tris buffered saline (TBS, 1x containing 1% BSA), and endogenous biotin with streptavidin/ biotin blocking kit (SP-2002, Vector Laboratories) for 15 min each at room temperature (RT). Slides were washed 3x with TBST (0.025% Triton X-100) for 1 min after each incubation. Tissue sections were incubated with primary antibodies (CD11b-5 µg/mL, #557395, BD Bioscience; Cox-1, 1 µg/mL, #160109, Cayman Chemicals; mPges-1-1.:200, #NBP1-87852, Novus Biologicals) in blocking solution overnight at 4ºC. Isotype-matched controls were included for non-specific binding of primary antibodies. Tissue sections were washed 3x with TBST before host -specific biotinylated-IgG secondary antibody (1 ug/mL in TBS+1% BSA, Vector Laboratories) were applied and incubated for 1 hour at RT. Peroxidase-conjugated Streptavidin (1 µg/mL, #016-030-084, Jackson ImmunoResearch) solution was then applied to tissue sections and incubated for 30 min at RT. Slides were immersed in MilliQ H2O for 5 min at RT, and then developed with ImmPACT DAB substrate kit (#SK-4105, Vector Laboratories) following manufacturer's instruction. Tissue slides were counterstained with hematoxylin, dehydrated and mounted in Permount mounting medium (#SP-15 100, Fisher Scientific).

**Aortic enzymatic dissociation and flow cytometry**

Mouse aortas (ascending aorta, aortic arch, descending aorta, innominate artery, carotid arteries, subclavian arteries, and descending aorta) were dissected and cut into ~1 mm pieces from the mPGES-1+/+/ Pd-1-/-/Ldlr-/- and mPges-1-/-/Pd-1-/-/Ldlr-/- (n = 4/group) for flow cytometry. Enzymatic tissue dissociation was conducted as previously described with some modifications (3). In brief, the aortic tissue fragments were immersed in 1.5 ml of enzyme cocktail containing DNase (120 U/ml; Worthington, #LS006331), Liberase™ (4 U/ml; Roche, #05401127001), and hyaluronidase (60 U/ml; Sigma-Aldrich, #H3506) in a petri dish and placed in an incubating shaker at 180 RPM at 37 °C for 45 min (VWR). To inactivate the enzymatic dissociation, the cells were filtered through a 70 µm strainer and washed with 10% fetal bovine serum (FBS, HyClone, #SH30071.03) in RPMI1640 (Gibco, #1187-085). The cells were centrifuged at 500 x g for five minutes at 4°C. The supernatant was decanted and the red blood cells were lysed by incubating the cell suspension with red blood cell lysing buffer (Hybri-Max; Sigma-Aldrich, #R7767) at room temperature for 1 min, which was also inactivated by 10% FBS in RPMI1640. The cells were washed two more times to remove debris with FACS buffer containing 5 mM EDTA (Invitrogen, #15575-038), 20 mM HEPES (Gibco, #15630-080), and 1mM sodium pyruvate (Gibco, #11360-070) in 1x PBS (Gibco, #14190-136). The washed cells were resuspended in FACS buffer for antibody staining. Briefly, ~1 x $10^6$ live cells were blocked with 1 µg of anti-CD16/32 antibody (TruStain TcX™, clone 93, #101320, BioLegend) for 10 min at room temperature. To this, staining buffer consisting of FACs and brilliant stain buffer (50 µL/test; Invitrogen, #00-4409-42) were added containing an assortment of conjugated antibodies outlined in supplementary Table X. The cells were stained with the antibody cocktail on ice in the dark for 20 min and resuspended in FACS buffer for flow cytometry analysis using the Cytek(R) Aurora equipped with the SpectroFlo3 software at the Children's Hospital of

Philadelphia Flow Cytometry Core. Data was analyzed using FCS Express$^{TM}$ (Version 7; De Novo Software) and represented using Prism.

**Table 1 :** Antibody cocktail.

| Antibodies | Manufacturer | Dilution | Ordering Number |
|---|---|---|---|
| CD45 PerCP/Cyanine5.5 (QA17A26) | Biolegend | 1:250 | 157612 |
| CD11b APC-Cyanine7 (M1/70) | Biolegend | 1:250 | 101226 |
| CD301 Alexa Fluor 647 (ER-MP23) | Novus Biologicals | 1:250 | NB100-64874 |
| CD3 FITC (17A2) | BD Biosciences | 1:250 | 555274 |
| F4/80 Brilliant Violet 650 (BM8) | Invitrogen | 1:250 | 416-4801-80 |
| CD11c eFluor 450 (N418) | Invitrogen | 1:250 | 48-0114-80 |
| Ly-6c PE (HK1.4) | Invitrogen | 1:3000 | 12-5932-80 |
| CD195 PE/Cyanine7 | Biolegend | 1:250 | 107017 |
| CD4 Brilliant Violet 615 (RM4-5) | Invitrogen | 1:250 | 366-0042-80 |
| CD8a Pacific Orange (5H10) | Invitrogen | 1:250 | MCD0830 |
| CD19 Brilliant Violet (eBio1D3) | Invitrogen | 1:250 | 417-0193-80 |
| CD9 Alexa Fluor 700 (eBioKMCB) | Invitrogen | 1:250 | 56-0091-82 |
| CD170 Brilliant Ultra Violet 737 (1RNM44N) | Invitrogen | 1:250 | 367-1702-80 |
| Ly-6g Brilliant Ultra Violet 395 (1A8-Ly6g) | Invitrogen | 1:230 | 363-9668-80 |
| Live/Dead Fixable Blue | Invitrogen | 1 µl/test | L34961 |

**Plasma analyses**

Commercially available testing kits were used for plasma analyses following manufacturer's instructions; glucose (Glucose Liqui-UV, #1060, Stanbio), Total cholesterol (Stanbio Cholesterol LiquiColor, #1010), HDL-C (Direct HDL-Cholesterol LiquiColor, #05902), and triglycerides (Stanbio LiquiColor Triglycerides, #2100), L-Type LDL-C (FujiFilm, #993-00404 & 999-00504). Lipoprotein lipase activity assay kit (Flourometric, #ab204721, abcam). Plasma non-HDL-C levels were calculated by subtracting LDL-C from total cholesterol levels.

**Preparation of bone-marrow derived macrophages (BMDM)**

Controls and EPr4$^{-/-}$ mice on standard laboratory diet (SLD) or WD for 12 weeks were used for the isolation of BMDM as described by Nguyen et al.(4). Briefly, mice euthanized by $CO_2$ exposure were sprayed with 70% ethanol before dissection. Skin and skeletal muscle were dissected to isolate femur and tibia from both legs. Bones with ends cut off were put into a sterile 0.5 ml reaction tube with a punched hole in the bottom, and 200µl of sterile RPMI cell culture medium was added into the tube. The reaction tube was transferred into a 1.5 mL collection tube, and bone marrow was collected in RPMI by spinning at 2600 g for 2 mins at room temperature. Red blood cells (RBCs) were lysed using Red Blood Cell Lysing Buffer Hybri-Max (R7767, Sigma). Bone marrow cells were cultured in RPMI for 5 days and differentiated in RPMI (70%)/ F929 conditioned medium (30%). BMDM was culture in DMEM (70%)/ F929 (30%) for chemotaxis assay. Ind-Cre$^{+/-}$/ Pd-1$^{-/-}$/ Ldlr$^{-/-}$ or EPr4$^{F/F}$/ Pd-1$^{-/-}$/Ldlr$^{-/-}$ (as control mice) and Ind-Cre$^{+/-}$/ EPr4$^{F/F}$/ Pd-1$^{-/-}$/ Ldlr$^{-/-}$ (as global EPr4$^{-/-}$ mice).

**Chemotaxis assay with bone-marrow derived macrophages (BMDM)**

300 µL of BMDM (0.5 x 10^6 cells) in DMEM/ F929 was seeded into a cell culture insert (8.0 µm pore size, #353097, Falcon) and placed in a 24-well plate containing 300 µL of DMEM/

F929 medium. After 24 hours, fresh medium was added, the plate was incubated for 24 hours in a $CO_2$ incubator at 37ºC.

Crystal violet was used to determine cell viability as described by Feoktistova et al.(5). Cell culture medium in the insert was aspirated, the insert was transferred into a clean 24-well plate with 500 µl of crystal violet solution (0.5%) for 20 min. The insert was rinsed 5x in distilled water, cells remained on the top of the insert were removed using a cotton-tipped swab (2x). The insert was air-dried for 6 hours, and crystal violet stain was dissolved with methanol on a rotation shaker for 15 min. 100 uL of dye mixture was transferred to a 96-well microtiter plate and optical density at 560 nm was measured using a plate reader.

**Cell viability assay of bone-marrow derived macrophages (BMDMs) using crystal violet dye**

BMDMs were loaded into 96-well plates at a density of 20,000 cells/ well in DMEM+F909 media for 6, 24, 48 and 72 hours. After incubation at 37ºC at each time point, cell culture media was removed, crystal violet solution (100 µL, 0.5% in water containing 20% methanol) was added into the wells for 20 mins at room temperature (RT). Wells were washed 4 times with distilled water and dried at RT for 4 hours. Crystal violet dye was dissolved with methanol (100 uL) at RT on a shaker for 15 mins. Optical density (OD) at 560 nm was measured using a plate reader. ODs at 24, 48 and 72 hours were normalized with ODs at 6 hours.

**Preparation of mouse aorta for real-time PCR analysis of gene expression**

Aortas from mice fed a WD were collected and snapped frozen with liquid nitrogen. All tissue samples were transferred to -80°C for storage until analyses. RNA was extracted using miRNeasy Mini Kit (Qiagen, Valencia, CA) following manufacturer's protocol. The concentration and quality of extracted RNA from platelets and MKs were measured using

NanoDrop® One (Thermo Scientific, Wilmington, DE) and reverse-transcribed into cDNA using Taqman Reverse Transcription Reagents (Applied Biosystems, Foster City, CA). Quantitative real time PCR was performed using Taqman Gene Expression Assays for Ptgs-1/ Cox-1 (Mm0125747_g1), Ptgs2/ Cox-2 (Mm00478374_m1), Ptges/ mPges-1 (Mm00452105_m1), EPr1 (00443097_m1), EPr2 (Mm00436051_m1), EPr3 (Mm001316856_m1), and EPr4 (Mm00436053_m1). An Applied Biosystems ViiA 7 real-time PCR system in a 384 well plate. Results were normalized to HPRT (Mm01545399_m1).

**Fluorescence activated cell sorting (FACS)**

To collect enough immune cells from mouse aortas (ascending aorta, aortic arch, descending aorta, innominate artery, carotid arteries, subclavian arteries, and descending aorta) for FACS, aortic cells were pooled from six mice per group after enzymatic dissociation to obtain enough immune cells for single cell sequencing. The aortic cells after dissociation as described above were resuspended in FACS buffer for antibody staining. Briefly, ~1 x $10^6$ live cells in FACS buffer (DPBS containing 2% FBS, 5mM EDTA, 20 mM HEPES, 1 mM sodium pyruvate) were blocked with 100 μL of anti-CD16/32 antibody (TruStain TcX™, clone 93, #101320, BioLegend) for 5 min at room temperature. The cells were stained with a PE/Cyanine7 α-mouse CD45 antibody (clone: 30-F11, #103114, BioLegend) in a 1:200 dilution on ice for 20 min after washing off blocking antibody and resuspended in FACS buffer for sorting. The cells were sorted using the Aurora CS cell sorter and SpectroFlo software (Cytekbio, Bethesda, MD, USA). The cells were collected in sheath fluid and transported to the single cell sequencing core for counting and downstream analyses.

**Single-cell RNA-sequencing of mouse aortas**

Male on $Ldlr^{-/-}$ mice after feeding a HFD for 24 weeks were euthanized, and aortas (ascending aorta, aortic arch, descending aorta, innominate artery, carotid arteries, subclavian arteries, and descending aorta) were dissected for single cells isolation with a cocktail of enzymes.[(6)] Briefly, dissected mouse aorta was cut into small pieces and incubated with 1.5 mL of enzyme cocktail (DNase- 120 U/mL, liberase- 4 U/mL, hyaluronidase- 60 U/mL) in a petri dish at 37ºC for 40 mins. A pipette was used to dissociate the tissue every 10 mins during the digestion. Cell supernatant was filtered through a 40 µm strainer and washed with RPMI1640 containing 10% FBS to inactive the enzyme cocktail. The cells were washed two more times to remove debris after performing RBC lysis and resuspended in DMEM/F12 media containing 10% FBS for further analysis. Mouse aortas from two mice per genotype were pooled after enzymatic digestion for scRNA- sequencing.

**scRNA-seq data analysis**

Sample demultiplexing, sequencing reads alignment to 10x Genomics prebuilt mouse reference transcriptome, cell calling, and gene expression quantification was performed with CellRanger (7), version 7.2.0. Downstream analysis was guided by the best practices initiative for single-cell data analysis (8) and executed in ScanPy (9, 10) framework, version 1.9.6. Apoptotic and low-quality cells were removed based on their gene expression and mitochondrial content. Detection of doublets was performed using scDblFinder (11), version 1.16.0. After filtering, 4232 cells in control and 9382 in $EPr4^{-/-}$ sample remained, with a median of 3541 and 2169 genes detected per cell, respectively. Counts were transformed and normalized using shifted logarithm with scran (12) (version 1.30.0) estimated size factors. Highly deviant genes were determined with scry (13) version 1.14.0. Clusters obtained using Leiden algorithm were annotated manually, based on the published list of marker genes specific to our experimental conditions (14) [Figure 7]

(Supplemental Figure S4). The manual annotation overall agreed with less specific annotation inferred by CellTypist (15) (version 1.6.2) low-resolution immune model. Annotated data were visualized with UMAP (16), version 0.5.5. Differential gene expression analysis was performed using PyDESeq2 (17) on the set of six pseduobulk samples, three per condition (control and $EPr4^{-/-}$). Pseudobulk samples were constructed by randomly splitting the collection of cells in each sample into three groups and then summing up gene counts in the cells belonging to individual groups. GSEA pathway analysis was done with decoupler (18) Python package, version 1.5.0. Ind-$Cre^{+/-}$/ Pd-$1^{-/-}$/ $Ldlr^{-/-}$ or $EPr4^{F/F}$/ Pd-$1^{-/-}$/$Ldlr^{-/-}$ (as control mice) and Ind-$Cre^{+/-}$/ $EPr4^{F/F}$/ Pd-$1^{-/-}$/ $Ldlr^{-/-}$ (as global $EPr4^{-/-}$ mice).

**Complete blood count analysis**

160 µL of whole blood from Pd-$1^{-/-}$/ $Ldlr^{-/-}$, mPges-$1^{-/-}$/ Pd-$1^{-/-}$/ $Ldlr^{-/-}$, controls and $EPr4^{-/-}$ mice on Pd-$1^{-/-}$/ $Ldlr^{-/-}$ background was collected via submandibular vein with a sterile lancet into a K2E K2EDTA coated tube. CBC was performed following manufacturer's instruction using Sysmex Hematology Analyzer (XN-V instrument). Ind-$Cre^{+/-}$/ Pd-$1^{-/-}$/ $Ldlr^{-/-}$ or $EPr4^{F/F}$/ Pd-$1^{-/-}$ /$Ldlr^{-/-}$ (as control mice) and Ind-$Cre^{+/-}$/ $EPr4^{F/F}$/ Pd-$1^{-/-}$/ $Ldlr^{-/-}$ (as global $EPr4^{-/-}$ mice).

**Mass spectrometric analysis of urinary prostaglandin metabolites**

Urinary prostanoid metabolites were measured by liquid chromatography / mass spectrometry as described (19). Such measurements provide a noninvasive, time integrated measurement of systemic prostanoid biosynthesis (20). Briefly, mouse urine samples were collected using metabolic cages over an eight hour period (9am to 5pm). Systemic production of $PGI_2$, $PGE_2$, $PGD_2$, and $TxA_2$ was determined by quantifying their major urinary metabolites- 2, 3-dinor 6-keto $PGF_{1\alpha}$ (PGIM), 7-hydroxy-5, 11-diketotetranorprostane-1, 16-dioic acid (PGEM), 11, 15-

dioxo-9$_{\alpha}$-hydroxy-2, 3, 4, 5-tetranorprostan-1, 20-dioic acid (tetranor PGDM) and 2, 3-dinor $TxB_2$ (TxM), respectively. Results were normalized with urinary creatinine.

**Mass spectrometric analysis of urinary creatinine**

Quantitation of plasma or urinary creatinine was performed using ultra high pressure liquid chromatography/tandem mass spectrometry (UPLC/MS/MS) with positive electrospray ionization and multiple reaction monitoring. A stable isotope-labeled internal standard (1mL for urine, 2.5 µg/mL [d3]-creatinine in 3% $H_2O$/acetonitrile) was added to 10 µL of mouse urine. The resulting solution of urine samples was further diluted by 10 times. Samples were transferred into an autosampler vial and 20 µL was injected to the UPLC-MS/MS system. A Shimadzu Prominence UPLC system was used for chromatography. The UPLC column was a 2.1 x 50 mm with 2.5 µm particles (Waters XBridge BEH HILIC). The mobile phase A was 100% acetonitrile. The mobile phase B was 5mM ammonium formate (pH = 3.98). The flow rate was 350 µL/min. Separations were carried out with fixed solvent gradients (12% mobile phase B). The Thermo Finnigan TSQ Quantum Ultra tandem instrument (Thermo Fisher Scientific) equipped with a triple quadrupole analyzer was operated in positive-mode ESI and the analyzer was set in the MRM mode for the analysis of creatinine. The transition for creatinine was 114>86. Quantitation was done by peak area ratio and results were normalized to the sample volume.
